# Xist-mediated silencing requires additive functions of SPEN and Polycomb together with differentiation-dependent recruitment of SmcHD1

**DOI:** 10.1101/2021.11.02.466758

**Authors:** Joseph S. Bowness, Tatyana B. Nesterova, Guifeng Wei, Lisa Rodermund, Mafalda Almeida, Heather Coker, Emma J. Carter, Artun Kadaster, Neil Brockdorff

**Affiliations:** Department of Biochemistry, University of Oxford, South Parks Road, Oxford, OX1 3QU, UK

## Abstract

X chromosome inactivation (XCI) is mediated by the non-coding RNA Xist which directs chromatin modification and gene silencing in *cis*. The RNA binding protein SPEN and associated corepressors have a central role in Xist-mediated gene silencing. Other silencing factors, notably the Polycomb system, have been reported to function downstream of SPEN. In recent work we found that SPEN has an additional role in correct localisation of Xist RNA in *cis*, indicating that its contribution to chromatin-mediated gene silencing needs to be reappraised. Making use of a SPEN separation-of-function mutation we show that SPEN and Polycomb pathways in fact function in parallel to establish gene silencing. Additionally, we find that differentiation-dependent recruitment of the chromosomal protein SmcHD1 is required for silencing many X-linked genes. Our results provide important insights into the mechanism of X inactivation and the coordination of chromatin-based gene regulation with cellular differentiation and development.

## Introduction

X chromosome inactivation (XCI) evolved in mammals to equalize the levels of X-linked gene expression in XX females relative to XY males (Lyon, 1961). The XCI process, which is developmentally regulated, is orchestrated by the X inactive specific transcript (Xist), a 17kb non-coding RNA, which accumulates in *cis* across the future inactive X (Xi) chromosome (Brockdorff et al., 1992; Brown et al., 1992; Lee and Jaenisch, 1997; Penny et al., 1996). Xist RNA recruits several factors that collectively modify chromatin/chromosome structure to silence X-linked genes (Boeren and Gribnau, 2021). Whilst XCI commences rapidly with the onset of Xist RNA expression, gene silencing is established progressively over a period of days (Borensztein et al., 2017; Lin et al., 2007; Marks et al., 2015; Sousa et al., 2019). Several X-linked genes exhibit either partial or complete escape from X inactivation (Posynick and Brown, 2019).

Gene silencing by Xist RNA is mediated principally by the A-repeat element, a tandem repeat located at the 5’ end of the transcript (Wutz et al., 2002). Recent studies have identified the RNA binding protein (RBP) SPEN as the critical silencing factor which recognises this element (Lu et al., 2016; McHugh et al., 2015; Monfort et al., 2015). Thus, SPEN loss of function and deletion of the A-repeat both result in strong abrogation of chromosome silencing (Chu et al., 2015; Dossin et al., 2020; McHugh et al., 2015; Moindrot et al., 2015; Monfort et al., 2015; Nesterova et al., 2019). SPEN interacts with the corepressor NCoR-HDAC3 through a C-terminal SPOC domain (Ariyoshi and Schwabe, 2003; Dossin et al., 2020), and this pathway plays an important role in Xist-mediated silencing (McHugh et al., 2015; Żylicz et al., 2019). Recent evidence indicates that the SPEN SPOC domain may also recruit other factors that contribute to Xi gene silencing (Dossin et al., 2020), and additionally that SPEN has a SPOC-independent function in ensuring localization and accumulation of Xist RNA across the Xi chromosome (Rodermund et al., 2021). Accordingly, deletion/mutation of the SPEN SPOC domain alone does not fully recapitulate the silencing deficiency observed in SPEN null cells or following deletion of the A-repeat (Dossin et al., 2020; Rodermund et al., 2021).

Several factors in addition to SPEN have been implicated in Xist-mediated silencing (reviewed in Brockdorff et al., 2020). The most notable example is the Polycomb system, comprising the multiprotein Polycomb repressive complexes PRC1 and PRC2 that catalyse the histone modifications H2AK119ub1 and H3K27me3 respectively (de Napoles et al., 2004; Plath et al., 2003; Silva et al., 2003). Polycomb recruitment to Xi is initiated by the RBP hnRNPK which also binds to a tandem repeat element in Xist RNA exon 1; the B/C-repeat (herein PID region) (Bousard et al., 2019; Colognori et al., 2019; Nesterova et al., 2019; Pintacuda et al., 2017). hnRNPK interacts with PCGF3/5-PRC1 complexes to promote deposition of H2AK119ub1. This step triggers a positive feedback cascade leading to recruitment of other PRC1 complexes and also PRC2 (Almeida et al., 2017). PCGF3/5 loss of function or deletion of the Xist PID region impairs Xi gene silencing, albeit not to the same degree as SPEN loss of function (Bousard et al., 2019; Colognori et al., 2019, 2020; Nesterova et al., 2019). Additionally, Polycomb-mediated gene silencing in XCI has been attributed principally to PRC1 and H2AK119ub1 rather than PRC2/H3K27me3 (Almeida et al., 2017; Nesterova et al., 2019). Similar to SPEN, the Polycomb pathway has been reported to have a secondary function in Xist RNA localization (Colognori et al., 2019).

Recruitment of both SPEN and the Polycomb system to Xi occurs rapidly at the onset of Xist RNA expression, consistent with their role in early establishment of gene silencing (Dossin et al., 2020; Nesterova et al., 2019; Żylicz et al., 2019). Conversely, some other XCI-associated factors, notably the histone variant macroH2A and the chromosomal protein SmcHD1, are concentrated on Xi after several days of Xist RNA expression, as determined by analysis of differentiating mouse XX embryonic stem cells (mESCs) (Gendrel et al., 2012; Mermoud et al., 1999). SmcHD1 recruitment to Xi is dependent on establishment of PRC1-mediated H2AK119ub1 (Jansz et al., 2018a). Loss of SmcHD1 function results in female-specific embryo lethality attributable to a failure of silencing in a subset of Xi genes (Blewitt et al., 2008; Gendrel et al., 2013; Mould et al., 2013). There is evidence that SmcHD1 facilitates long-term maintenance of X inactivation (Sakakibara et al., 2018), but a possible role in establishment of gene silencing has not been investigated in depth. Additionally, SmcHD1 is important for DNA methylation of Xi CpG islands (Blewitt et al., 2008; Gendrel et al., 2012), and the establishment of a unique chromosome architecture on Xi (Gdula et al., 2019; Jansz et al., 2018b; Wang et al., 2018).

In this study we analyse a series of novel loss of function mESC lines to further elucidate the interplay of SPEN, Polycomb and SmcHD1 in Xist-mediated X chromosome silencing.

## Results

### Defining the role of the SPEN SPOC domain in Xist-mediated gene silencing

In recent work we derived SPEN^SPOCmut^ interspecific XX mESC lines with point mutations that abrogate SMRT/NCoR interaction with the SPEN-SPOC domain (Ariyoshi and Schwabe, 2003; Oswald et al., 2016; Rodermund et al., 2021), and found that Xist-mediated silencing is only partly reduced, contrasting with reports using complete loss of function SPEN mutations (Dossin et al., 2020; Monfort et al., 2015; Nesterova et al., 2019). Equivalent conclusions were reached in an independent study that analysed XX mESCs with a deletion of the SPOC domain (Dossin et al., 2020). Indeed, the degree of silencing observed after 1 day of Xist induction is highly similar in the two studies (Figure S1A), suggesting that the precise SPEN^SPOCmut^ point mutation has an equivalent biological effect to that seen with complete deletion of the SPOC domain. The retained silencing activity following mutation of the SPOC domain is referred to henceforth as SPOC-independent silencing.

To further investigate SPOC-independent silencing we extended our analysis across a more complete X inactivation timecourse. Thus, we optimized a protocol for deriving neuronal precursor cells (NPC) from mESCs (Conti et al., 2005; Splinter et al., 2011), enabling analysis of the temporal trajectory of Xist-mediated silencing in a highly synchronous and homogeneous model. We assayed X-linked gene silencing by sequencing Chromatin-associated RNA (ChrRNA-seq) over daily timepoints of NPC differentiation and thus found that Xist-mediated gene silencing is largely complete by around day 7 in wild-type (WT) interspecific XX lines with doxycycline-inducible Xist on the *M.m.domesticus* allele (iXist-ChrX_Dom_) (Figure S1B). Accumulation of Xist RNA, as quantified from ChrRNA-seq data, increased progressively over the same timecourse (Figure S1B).

We then analysed Xist-mediated silencing during NPC differentiation in SPEN^SPOCmut^ mESCs, which were derived from the iXist-ChrX_Dom_ parent line. Failure of X inactivation results in significant levels of X chromosome elimination in late stage differentiated XX mESCs, largely reflecting selection against cells with two active X chromosomes (Colognori et al., 2020), and we therefore limited our analysis to the first six days of differentiation. Consistent with our previous study (Rodermund et al., 2021), partial silencing is observed in SPEN^SPOCmut^ mESCs after 1 day of Xist induction (Figure 1A, Figure S1C). SPOC-independent silencing persisted and in fact marginally increased at later differentiation timepoints, up to day 6 of NPC differentiation (Figure 1A). These findings contrast with the near complete loss of silencing observed using the SPEN^ΔRRM^ mutation (Nesterova et al., 2019), and A-repeat deletion (Coker et al., 2020), as shown in Figure S1C. Levels of Xist RNA were slightly reduced compared to WT iXist-ChrX*_Dom_* (Figure 1B), but not to the dramatic extent observed in SPEN^ΔRRM^ or Xist^ΔA^ lines (Figure S1C). Downregulation of pluripotency markers (*Nanog* and *Klf2*) were similar in WT and SPEN^SPOCmut^ cells, although upregulation of neuron-specific markers (*Nes* and *Vim*) appeared deficient by differentiation day 6 in the latter (Figure S1D).

**Figure 1.**
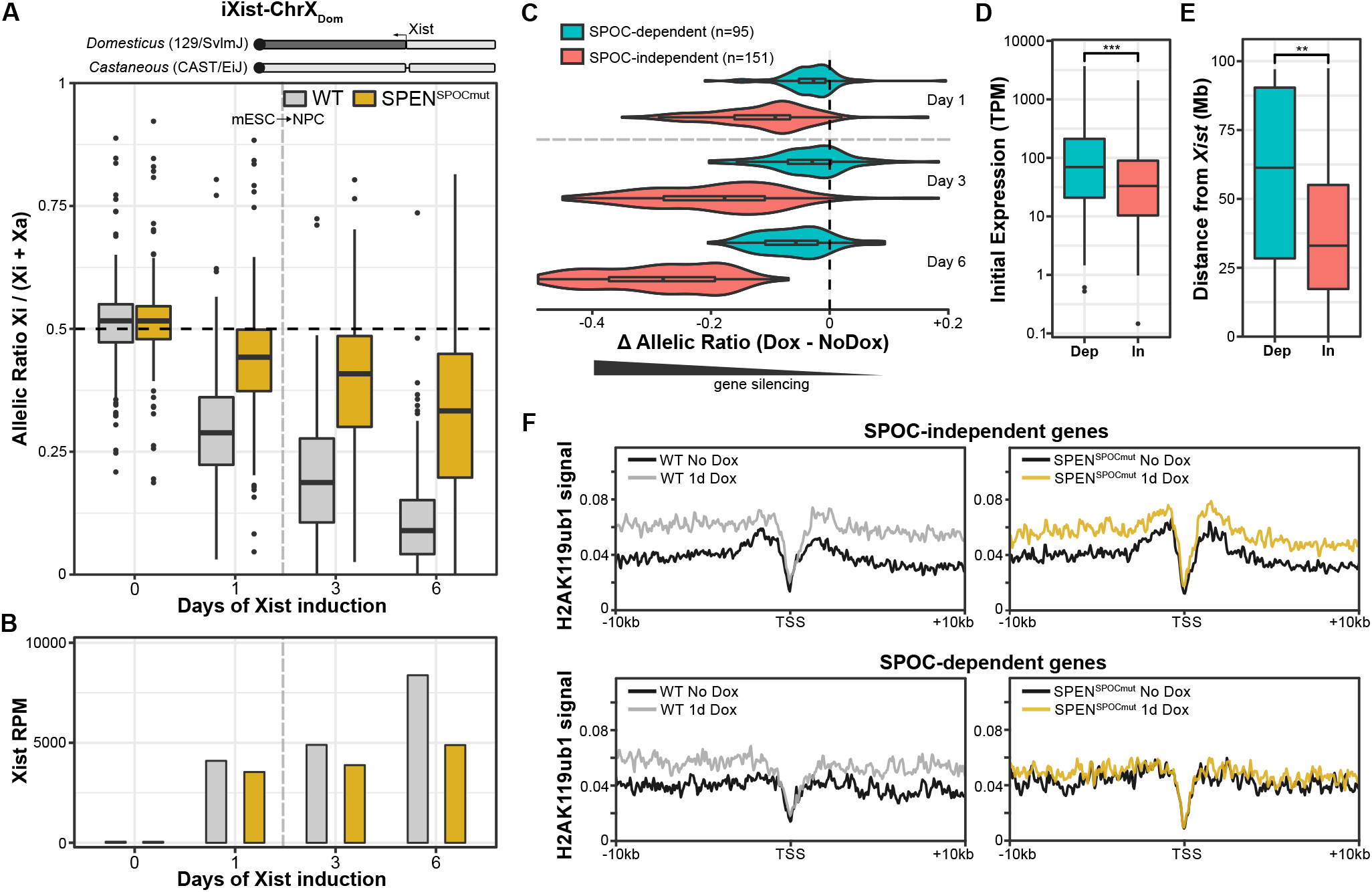
Xist-mediated silencing in SPEN^SPOCmut^ mESCs (A) (above) Schematic of the iXist-ChrX_Dom_ interspecific model cell line, described in detail in our previous study (Nesterova et al., 2019). Sequence originating from *Domesticus*/129 and *Castaneous* chromosomes are shown by dark and light shading respectively. (below) Boxplots summarizing allelic ChrRNA-seq analysis of X-linked gene expression in iXist-ChrX_Dom_ WT and SPEN^SPOCmut^ cell lines (n=246 genes). Timepoints in mESCs with and without Xist induction (day 1 and day 0 respectively) are separated from timepoints of Xist induction under NPC differentiation conditions (days 3 and 6) by the vertical dashed line. SPEN^SPOCmut^ ES boxes are averaged from the three clones presented in Figure S1C. Day 3 and day 6 NPC boxes are each averaged from duplicate experiments differentiating iXist-ChrX_Dom_ WT and SPEN^SPOCmut^ in parallel (clones D9 and H1). The horizontal dashed line at Allelic Ratio=0.5 represents equal expression of both alleles. (B) Relative levels of chromatin-associated Xist RNA for (A). (C) Violin plots of the gene silencing in SPEN^SPOCmut^ cells after 1, 3 and 6 days of Xist induction, as measured by the change in allelic ratio (Dox – NoDox). SPOC-independent and SPOC-dependent genes (see Figure S1E) are separated for comparison. Replicates for each timepoint are averaged together. (D) Boxplots comparing the initial expression levels of SPOC-dependent and SPOC-independent genes in iXist-ChrX_Dom_ mESCs (mRNA-seq data; GSEXXXX). (E) Boxplots comparing the genomic distance from the *Xist* locus of SPOC-dependent and SPOC-independent genes. (F) TSS-centred meta-profiles comparing enrichment of H2AK119ub1 for uninduced and 1 day-induced WT versus SPEN^SPOCmut^ mESCs at SPOC-dependent and SPOC-independent genes. Significance calculated by Wilcoxon signed-rank test (** indicates p *<*0.01, *** indicates p *<* 0.001).

To investigate the basis for SPOC-independent silencing we defined a gene subset that shows greater silencing in SPEN^SPOCmut^ cells after 6 days of Xist induction (Figure 1C, Figure S1E), and then compared their characteristics to those of other minimally-silenced, strictly SPOC-dependent, Xi genes. Using this approach we found that SPOC-independent silencing is associated with lower expressed genes and with genes that are closer to the *Xist* locus (Figure 1D,E). Additionally, we observed an association with a local chromatin environment enriched in H3K27me3 as defined by the ChromHMM model (Figure S1F) (Ernst and Kellis, 2017; Nesterova et al., 2019). Interestingly, these characteristics resemble those identified previously as being associated with Xi genes that are more affected by disruption of the Polycomb pathway (Nesterova et al., 2019; Pintacuda et al., 2017), indicating that the Polycomb system may underpin SPOC-independent silencing. Consistent with this idea, a re-examination of H2AK119ub1 native ChIP-seq from SPEN^SPOCmut^ (Rodermund et al., 2021) revealed a greater enrichment of H2AK119ub1 over SPOC-independent genes compared to SPOC-dependent genes after 24 hours of Xist induction in mESCs (Figure 1F).

### SPEN and Polycomb function additively to establish Xist-mediated silencing

To further investigate the link between SPOC-independent silencing and the Polycomb system we examined the effect of depleting Polycomb function in SPEN^SPOCmut^ mESCs. Accordingly, we made use of the FKBP12^F36V^/dTAG-13 degron system (Nabet et al., 2018) to acutely deplete PCGF3/5-PRC1, the complex required to initiate Polycomb recruitment to Xi (Almeida et al., 2017). Using CRISPR/Cas9-facilitated homologous recombination, we introduced an FKBP12^F36V^ degron to the N-termini of both PCGF3 and PCGF5 in iXist-ChrX_Dom_ mESCs. Treatment of the tagged mESC line, iXist-ChrX_Dom_ FKBP12^F36V^-PCGF3/5, with the cell-permeable small molecule dTAG-13 resulted in rapid and complete degradation of both proteins within 15-30mins with no detectable effect on protein levels of RING1B or SUZ12, core subunits of PRC1 and PRC2 respectively (Figure 2A). Consistent with a previous analysis of PCGF3/5 conditional knockout mESCs (Fursova et al., 2019), we observed a ∼30% global reduction in H2AK119ub1 determined by calibrated native ChIP-seq after 36 hours of dTAG-13 treatment, attributable to reduced ‘blanket’ coverage over intergenic or gene body regions rather than at canonical PRC1 target sites (Figure S2A). Genome-wide levels of H3K27me3 were broadly unchanged (Figure S2B).

**Figure 2.**
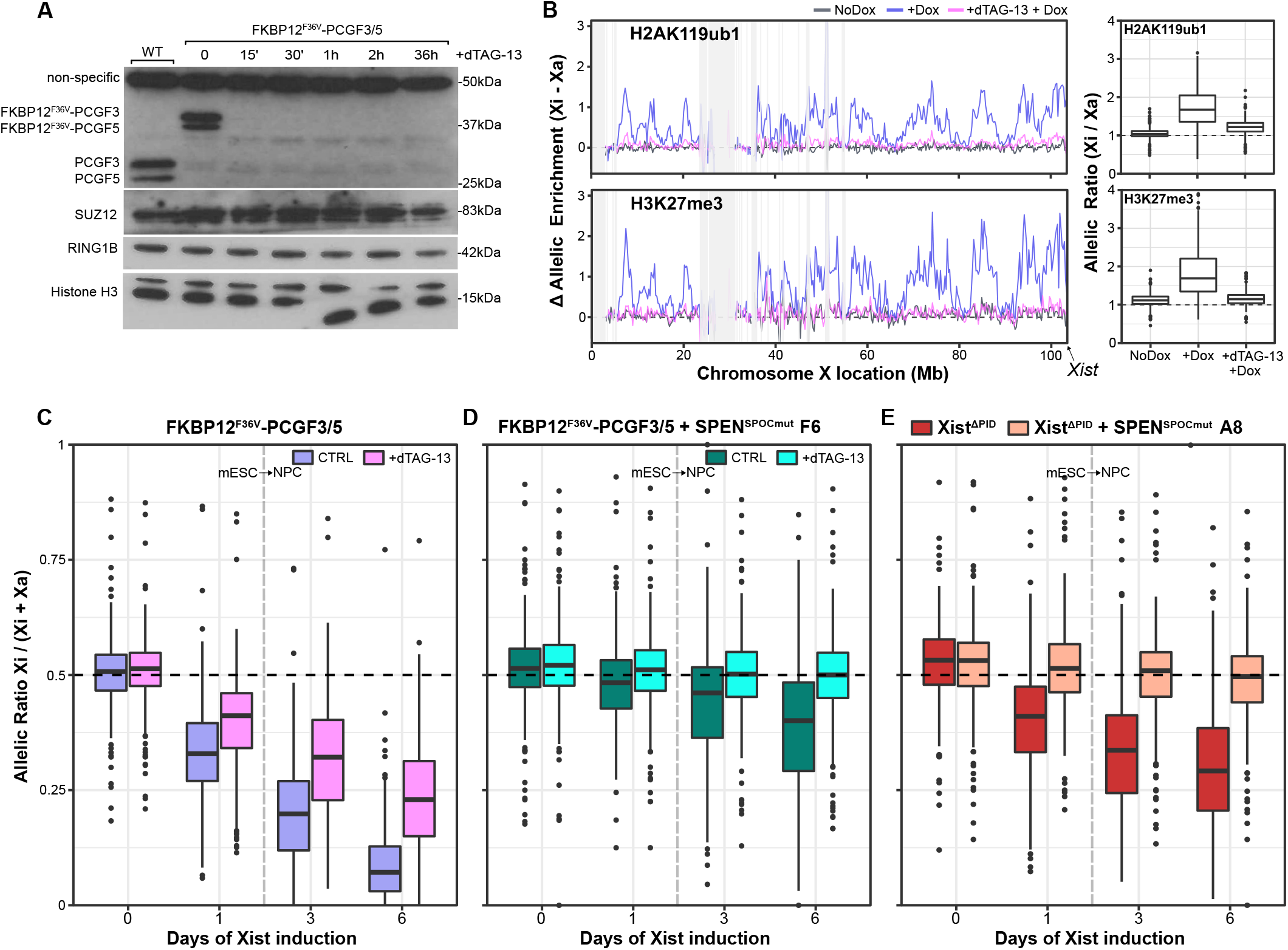
SPOC-independent silencing is attributable to the Polycomb system (A) Western blot showing rapid degradation of FKBP12^F36V^-PCGF3 and FKBP12^F36V^-PCGF5 fusion proteins within 15 minutes of dTAG-13 treatment. Histone H3 provides a loading control. (B) Plots showing allelic H2AK119ub1 (above) and H3K27me3 (below) enrichment over 250Kb windows after 1 day of Xist induction in untreated FKBP12^F36V^-PCGF3/5 cells and cells treated for 12 hours with dTAG-13 prior to Dox addition. Lines are averages of two highly correlated replicate experiments. Shaded regions mask blacklisted windows with low allelic mappability (see Methods). Boxplot quantification of allelic ratios (here calculated as Xi / Xa) for non-blacklisted 250kb windows from the line graphs (n = 335) are shown to the right. (C) Boxplots summarizing allelic ChrRNA-seq analysis of X-linked gene expression in the iXist-ChrX_Dom_ FKBP12^F36V^-PCGF3/5 cell line (n=246 genes). Timepoints in mESCs with and without Xist induction (day 1 and day 0 respectively) are separated from timepoints of Xist induction under NPC differentiation conditions (days 3 and 6) by the vertical dashed line. Untreated cells are compared to cells treated for 12 hours with dTAG-13 prior to Xist induction, cultured in parallel. All boxes show averages of two highly similar replicate experiments. (D) Boxplots summarizing allelic ChrRNA-seq analysis of X-linked gene expression in iXist-ChrX_Dom_ FKBP12^F36V^-PCGF3/5 + SPEN^SPOCmut^ cells for timepoints in mESCs with and without Xist induction (day 1 and day 0 respectively), and 3 and 6 days under NPC differentiation conditions (n=246 genes). Untreated cells are compared to cells treated for 12 hours with dTAG-13 prior to Xist induction, cultured in parallel. A replicate experiment from a separate clone is shown in Figure S2D. (E) As (D) for iXist-ChrX_Dom_ Xist^.6.PID^ + SPEN^SPOCmut^. Day 3 and 6 Xist^.6.PID^ samples are averages of duplicate experiments in which the parental Xist^.6.PID^ line was differentiated in parallel with one of two iXist-ChrX_Dom_ Xist^.6.PID^ + SPEN^SPOCmut^ derivative clones. This second clone is shown in Figure S2F. Horizontal dashed lines at Allelic Ratio=0.5 represent equal expression of both alleles (C-E).

iXist-ChrX_Dom_ FKBP12^F36V^-PCGF3/5 was further validated to confirm near-complete loss of Xi specific accumulation of Polycomb-mediated H2AK119ub1 and H3K27me3 as determined by allelic ChIP-seq analysis (Figure 2B). ChrRNA-seq of Xist-mediated silencing in cells treated with dTAG-13 for 12 hours prior to Xist induction demonstrated a moderate silencing deficiency (Figure 2C), equivalent to that reported in our previous work (Nesterova et al., 2019). Levels of Xist RNA were broadly similar between untreated and dTAG-13-treated samples (Figure S2C).

We went on to engineer the SPEN SPOC mutation in iXist-ChrX_Dom_ FKBP12^F36V^-PCGF3/5 mESCs to analyse the effect of PCGF3/5 degradation on SPOC-independent silencing. Strikingly, treatment of the combined mutants with dTAG-13 resulted in a complete loss of Xist-mediated silencing throughout NPC differentiation in two independent clones (Figure 2D, Figure S2D). As above, we only analysed differentiation timepoints up to day 6 due to progressive selection for X chromosome elimination in differentiated cells with two active X chromosomes. Levels of Xist RNA were equivalent to SPEN^SPOCmut^ clones (Figure S2E, cf. Figure 1B).

PCGF3/5-PRC1 complexes function globally in genome regulation and it is conceivable that indirect effects of perturbing other pathways contributes to the silencing deficit that we observed. Therefore, to further validate our findings we introduced the SPEN SPOC mutation into the iXist-ChrX_Dom_ Xist^ΔPID^ XX mESC line in which the Xist B/C-repeat region required for hnRNPK/Polycomb recruitment in X inactivation is deleted (Nesterova et al., 2019). Allelic ChrRNA-seq analysis was carried out to assess Xist-mediated gene silencing following Xist induction and NPC differentiation. As shown in Figure 2E and Figure S2F, SPOC-independent silencing was abolished in two independent clones. Levels of Xist RNA were similar to those seen in the parental Xist^ΔPID^ line (Figure S2G). Together these results further support the conclusion that SPOC-independent silencing in XCI is attributable to activity of the Polycomb system.

Abrogation of Xi Polycomb recruitment and the SPEN^SPOC^ mutation have both been reported to have effects on Xist RNA localization (Colognori et al., 2019; Rodermund et al., 2021), and this could potentially contribute to the aberrant silencing observed when both pathways are depleted. Qualitative analysis of Xist RNA domains using RNA FISH and conventional microscopy indicates monoallelic Xist upregulation and cloud formation after 1 day of Xist induction in mESCs for all cell lines presented hitherto (Figure S3A-D). We did however observe a tendency towards larger clouds in FKBP12^F36V^-PCGF3/5 cells after dTAG-13 treatment (Figure S3B), and occasional Xist RNA dispersal in double mutant cells (Figure S3C,D). With this in mind, we went on to quantify Xist localization parameters using computational analysis of super-resolution 3D-structured illumination microscopy (3D-SIM) of Xist RNA FISH in the iXist-ChrX_Dom_ FKBP12^F36V^-PCGF3/5 model. As summarized in Figure 3A-D, PCGF3/5 degradation alone and in combination with SPEN^SPOCmut^ resulted in expanded Xist RNA territories and an increased number of Xist RNA foci. Additionally, a proportion of cells had either partially or extensively dispersed Xist RNA, the latter category being larger with the combined mutation (Figure 3C,D). We conclude that mis-localization of Xist RNA may make some contribution to the observed gene silencing deficiencies, but given that Xist expression and Xi domain formation is maintained in most cells, the deficit is unlikely to account for the complete abolition of silencing in the combined knockout.

**Figure 3.**
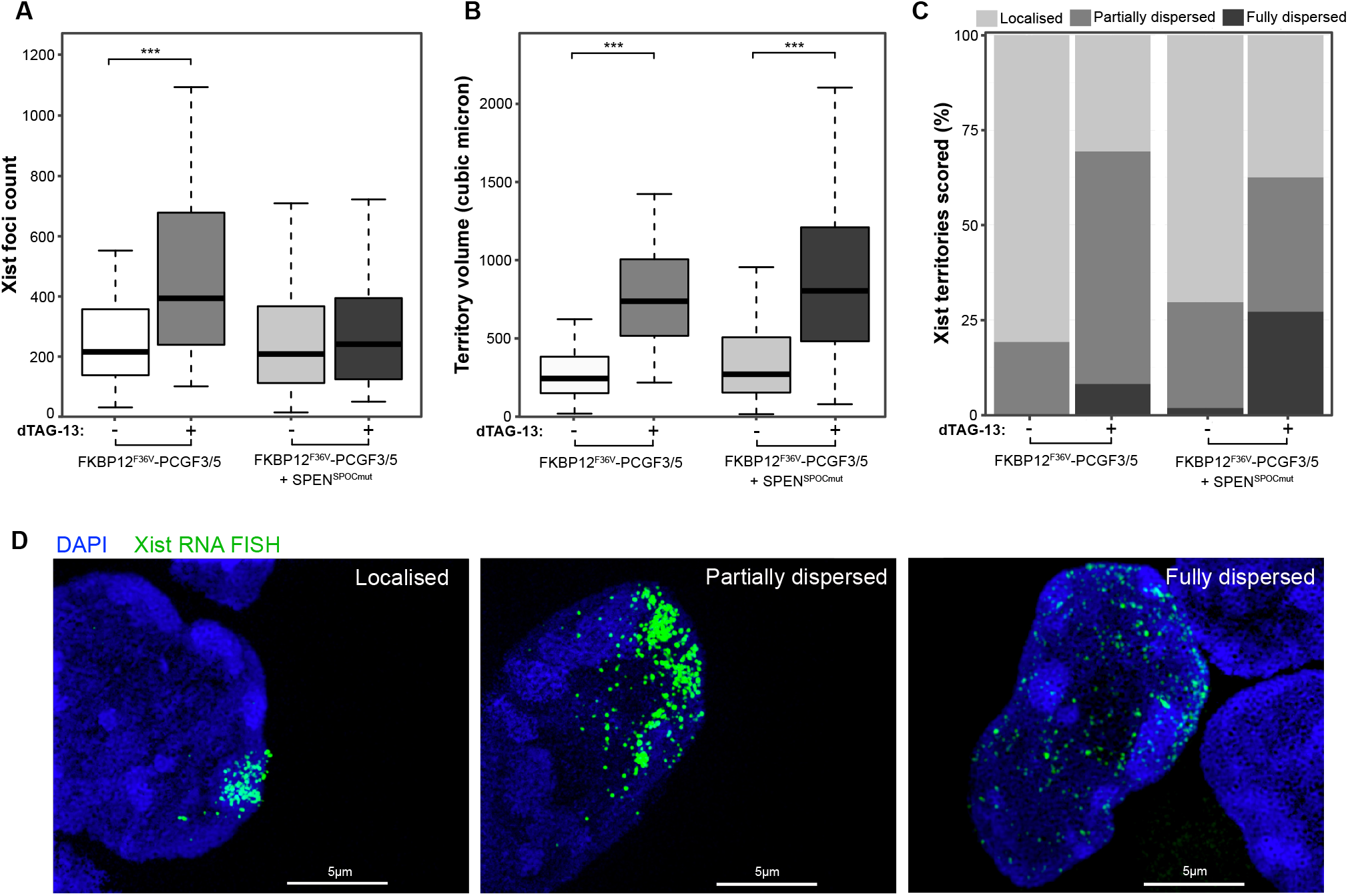
Quantitative 3D-SIM analysis of Xist RNA localization after PCGF3/5 degradation (A,B) Boxplots showing quantification of number of individual Xist foci (A) and Xist territory volume (B) in FKBP12^F36V^-PCGF3/5 and FKBP12^F36V^-PCGF3/5 + SPEN^SPOCmut^ cells. Comparisons are between mESCs after 1 day of induction with or without dTAG-13 treatment for 12 hours to induce degradation of PCGF3/5 prior to Xist expression. n = 40 cells per sample. *** indicates p*<*0.001 by unpaired two sample Wilcoxon test. (C) Scoring by eye of the proportions of cells in each sample with localized, partially dispersed, and fully dispersed Xist territories. n = 40 cells per sample. (D) Representative 3D-SIM images (z-projections) of cells scored in each category of Xist dispersal.

To summarize, our findings suggest that SPOC-independent silencing is in large part mediated by the Polycomb pathway, implying that Polycomb functions in parallel with rather than downstream of SPEN as the two principal pathways mediating the establishment of Xist-mediated gene silencing.

### Xist-mediated silencing is linked to cellular differentiation

Both SPEN and Polycomb systems are recruited to Xi rapidly at the onset of Xist RNA expression but silencing of individual genes proceeds over several hours and in some cases days (Borensztein et al., 2017; Lin et al., 2007; Loda et al., 2017; Marks et al., 2015; Nesterova et al., 2019; Sousa et al., 2019). Additionally, there is evidence that cellular differentiation promotes Xist-mediated silencing, albeit based on analysis of only two X-linked genes (Loda et al., 2017). To further address this latter point we assayed allelic silencing chromosome-wide after long-term (10 days) continuous Xist RNA induction in undifferentiated mESCs compared with NPC differentiation conditions. For these experiments we made use of both iXist-ChrX_Dom_ and a reciprocal interspecific XX mESC line with the inducible promoter driving Xist RNA expression on the *M.m.castaneous* X chromosome (iXist-ChrX_Cast_) (Figure 4A-C, Figure S4A,B). Unlike iXist-ChrX_Dom_, iXist-ChrX_Cast_ is informative across the entire X chromosome, enabling allelic expression analysis of a larger number of X-linked genes. As shown in Figure 4A,B, silencing is significantly reduced in undifferentiated mESCs compared to cells differentiated to NPCs over an equivalent timecourse. Principle component analysis of autosomal gene expression (Figure 4C), and marker gene analysis (Figure S4A,B) confirm retention of mESC identity in these experiments.

**Figure 4.**
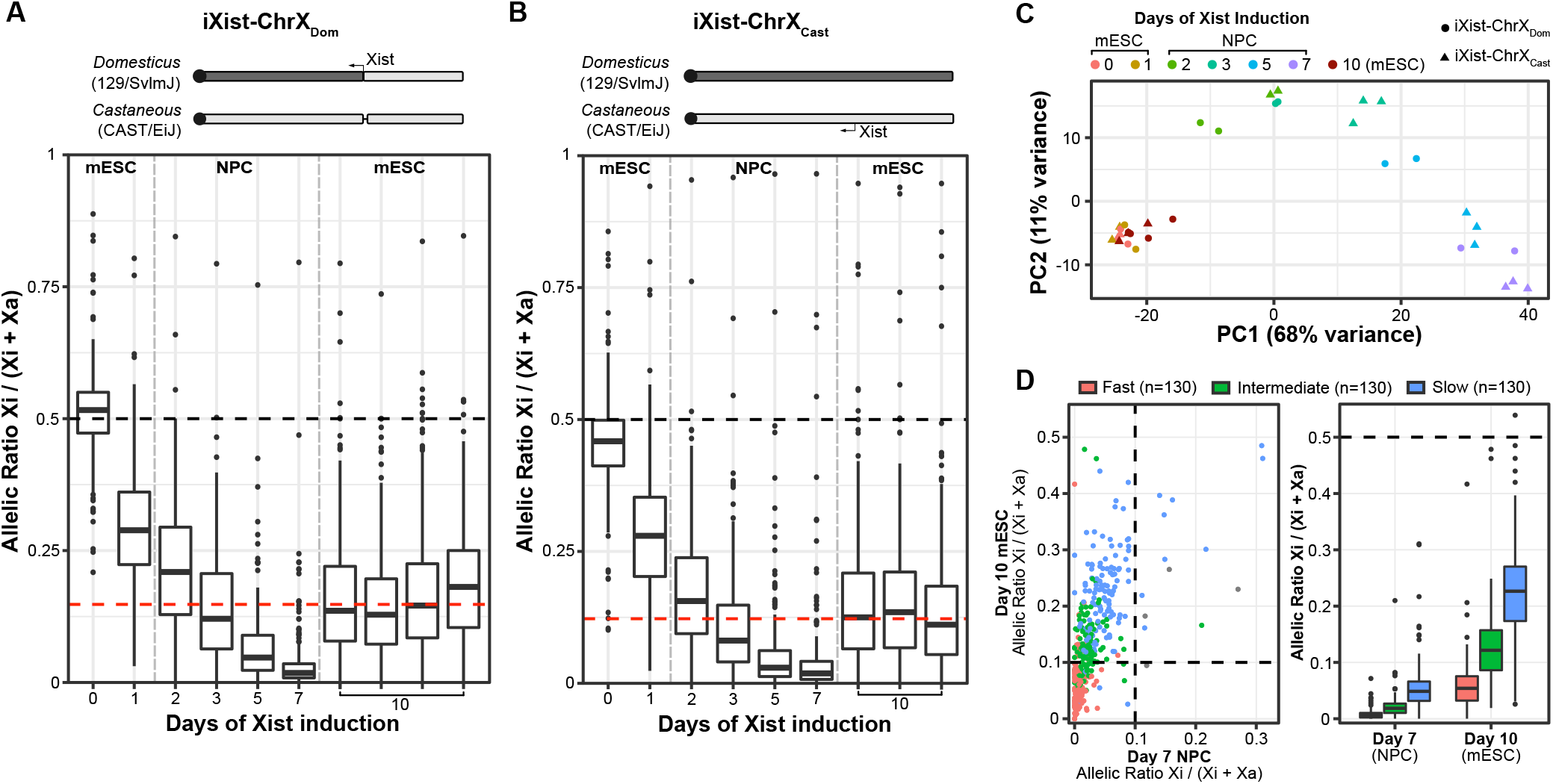
Cellular differentiation is required for completion of Xist-mediated silencing (A) Boxplots summarizing ChrRNA-seq analysis of X-linked gene expression in iXist-ChrX_Dom_ mESCs with 10 days of Xist induction in comparison to a 7-day timecourse of Xist induction under NPC differentiation conditions (n=246 genes). Four replicate experiments are displayed for undifferentiated mESCs, with the red horizontal dashed line indicating the median allelic ratio. A schematic of the iXist-ChrX_Dom_ model cell line is displayed above. See Figure 1A legend for further details. (B) As in (A) except for iXist-ChrX_Cast_ mESCs with 10 days of Xist induction in comparison to a timecourse of Xist induction under NPC differentiation conditions (n=399 genes). Three replicates of iXist-ChrX_Cast_ mESCs induced for 10 days are shown separately. NPC day 3, 5 and 7 boxes are averaged from triplicate samples. ES and NPC day 2 boxes are averaged from duplicate samples. A schematic of the iXist-ChrX_Cast_ cell line, in which Xist on the *M.m.Castaneous* allele is under doxycyline-inducible control, is displayed above. Horizontal dashed lines at Allelic Ratio=0.5 represent equal expression of both alleles, whilst vertical dashed lines delineate mESC versus NPC differentiation cell culture conditions (C-E). (C) PCA plot of each individual sample in (A) and (B) according to two leading principal components, defined from analysis of the top 500 differentially expressed autosomal genes across the whole data set. Colours indicate timepoints and shapes the model cell line. Day 10-induced mESCs cluster with day 0 and 1 mESCs, indicating that they retain embryonic stem state. (D) Scatter plot (left) comparing individual gene allelic ratios between iXist-ChrX_Cast_ mESCs with 10 days of Xist induction and NPC differentiation day 7. Dots represent individual genes, which are coloured according to the kinetic classes defined by fitting exponential models to silencing trajectories of NPC differentiation (see Figure S4C,E). Dashed lines trace allelic ratios of 0.1. Boxplots (right) compare mESCs with 10 days of Xist induction and NPC differentiation day 7. Genes are separated by kinetic silencing classes. The horizontal dashed line at Allelic Ratio=0.5 represents equal expression of both alleles.

We went on to determine if there is a relationship between dependence on cellular differentiation and silencing dynamics. Thus, we defined a silencing halftime (t_1/2_) for each X-linked gene by fitting the silencing trajectory with an exponential decay model, summarising over multiple ChrRNA-seq timepoints and replicates from NPC differentiation timecourses of iXist-ChrX_Dom_ and iXist-ChrX_Cast_ (Figure S4C). Halftimes were strongly correlated between cell lines (Figure S4D, R=0.82 Spearman’s Rank Correlation) and were used to classify genes into three equal sized groups showing fast, intermediate, and slow silencing kinetics (Figure S4E). Consistent with prior reports (Marks et al., 2015; Nesterova et al., 2019; Sousa et al., 2019), we found that proximity to the *Xist* locus and expression levels prior to Xist induction both influence dynamics of gene silencing, with higher-expressed genes and genes further from *Xist* silencing more slowly (Figure S4F). We then examined the relationship between silencing dynamics and the genes which fail to complete silencing following 10 days of Xist induction in undifferentiated mESCs, and found a clear correspondence between X-linked genes showing differentiation-dependence and intermediate/slow silencing groups (Figure 4D).

### Differentiation-linked Xi silencing correlates with SmcHD1 dependence

A possible explanation for the link between Xi silencing and differentiation is the involvement of a factor(s) not available in undifferentiated mESCs. A candidate for this function is the chromosomal protein SmcHD1, which is recruited to Xi dependent on PRC1 activity (Jansz et al., 2018a), but only after several days, based on analysis of XX mESCs undergoing embryoid body differentiation (Gendrel et al., 2012). Re-examination of the kinetics of SmcHD1 association with Xi by IF in our NPC model with inducible Xist expression confirms SmcHD1 recruitment is a late step, becoming detectable only after 3-4 days of Xist induction and NPC differentiation Figure 5A, B. Differences in timing relative to prior analysis using embryoid body differentiation may reflect the lag in onset of Xist RNA expression from the native vs. inducible promoter. Western blot analysis demonstrates that overall levels of SmcHD1 are equivalent in mESCs and throughout NPC differentiation (Figure 5C, Figure S5A), indicating that SmcHD1 availability of itself does not account for delayed Xi recruitment. SmcHD1 recruitment to Xi could not be detected in XX mESCs following 10 days of continuous induction of Xist RNA expression (Figure 5D,E).

**Figure 5.**
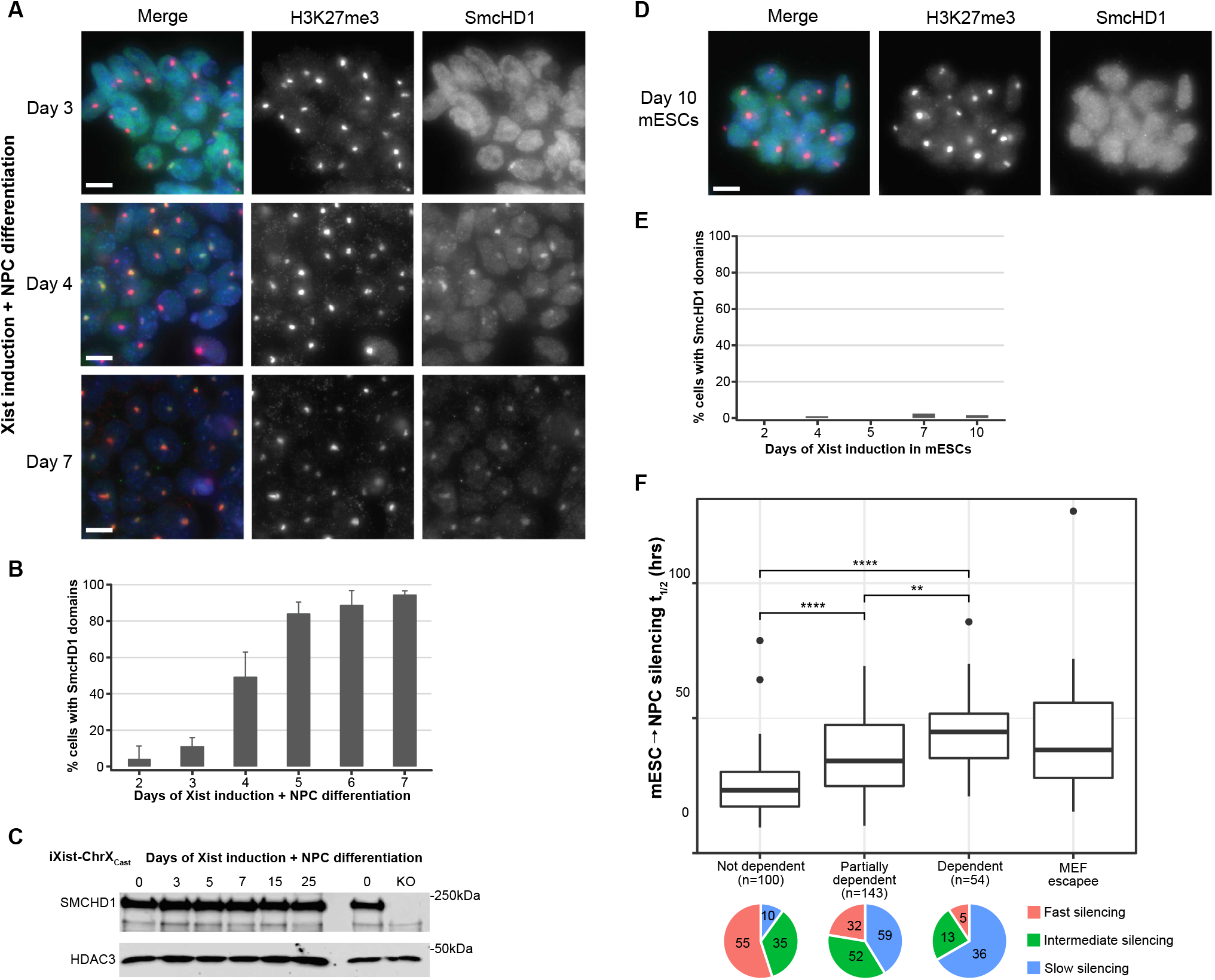
SmcHD1 recruitment to Xi is linked to differentiation-dependent silencing A) Immunofluorescence analysis of SmcHD1 and H3K27me3 (marker for presence of Xi) in the iXist-ChrX_Cast_ line during the timecourse of Xist induction under NPC differentiation conditions. Representative examples of cells at days 3, 4 and 7 of differentiation are shown. H3K27me3 is shown in magenta, SmcHD1 in cyan, and DNA in blue (DAPI) in the merged images. Scale bar is 10 *µ*m. B) Scoring of the number of cells showing SmcHD1 recruitment to Xi in the NPC differentiation timecourse. C) Western blot analysis in iXist-ChrX_Cast_ cells demonstrating that SmcHD1 protein levels are comparable in mESCs and during the NPC differentiation timecourse. A KO control is included to indicate the size of the specific SmcHD1 band. HDAC3 acts as a loading control. Molecular weight markers are shown on the right. D) SmcHD1 enrichment on Xi is not detectable after 10 days of Xist induction under ES conditions. Representative example images of mESCs induced with doxycycline for 10 days are shown. Scale bar is 10 *µ*m. E) Scoring of SmcHD1 domains in mESCs after 10 days of Xist induction. F) Boxplots comparing silencing halftimes between subsets of genes based on dependence on SmcHD1 for gene silencing in MEFs. Significance of individual comparisons is determined by Welch’s unequal variances T-test. ** and **** indicate p values below 0.01 and 0.0001 respectively. Pie charts below illustrate the proportions of genes within each SmcHD1-dependence group which have slow, intermediate, or fast kinetics of silencing in WT iXist-ChrX_Cast_ cells.

Differentiation-dependent recruitment of SmcHD1 was previously interpreted to support a role in maintenance rather than establishment of Xi gene silencing (Gendrel et al., 2012). However, silencing of genes showing intermediate/slow kinetics is incomplete at the time of SmcHD1 recruitment (Figure S5B), raising the possibility that SmcHD1 has a role in establishing silencing for these gene groups. With this in mind we performed an analysis of the relationship between silencing dynamics and SmcHD1 dependency, the latter being based on classifying genes as dependent (n=56), partially dependent (n=143), not dependent (n=101) or escapees (genes also expressed from Xi in WT, n=18) using ChrRNA-seq data from XX mouse embryo fibroblast (MEF) lines derived from SmcHD1 null embryos (Figure S5C and Gdula et al., 2019). As illustrated in Figure 5F we observed a clear association between SmcHD1 dependence and silencing dynamics, with SmcHD1-dependent genes strongly overlapping with slow silencing genes. Additionally, we found that SmcHD1 dependency is associated with genes showing incomplete silencing after long-term Xist expression in undifferentiated mESCs, illustrated in Figure S5D for the iXist-ChrX_Cast_ cell line.

### SmcHD1 facilitates differentiation-dependent silencing

The aforementioned observations suggest that SmcHD1 could have a role in establishment of Xi silencing (in addition to its defined role in maintenance of Xi silencing), specifically in relation to genes that exhibit a relatively slow silencing trajectory. To test this idea directly, we used CRISPR/Cas9 mediated mutagenesis to generate SmcHD1 knockout (KO) cell lines in iXist-ChrX_Dom_ and iXist-ChrX_Cast_ backgrounds (Figure S6A). Generation of homozygous KO clones was confirmed by western blot analysis (Figure 6A). Knockout lines were then validated to confirm that enrichment of Polycomb-linked histone modifications over Xi is maintained (Figure S6B), and furthermore that nuclear SmcHD1 signal and Xi enrichment is absent after 7 days of Xist induction under NPC differentiation conditions (Figure 6B, Figure S6C).

**Figure 6.**
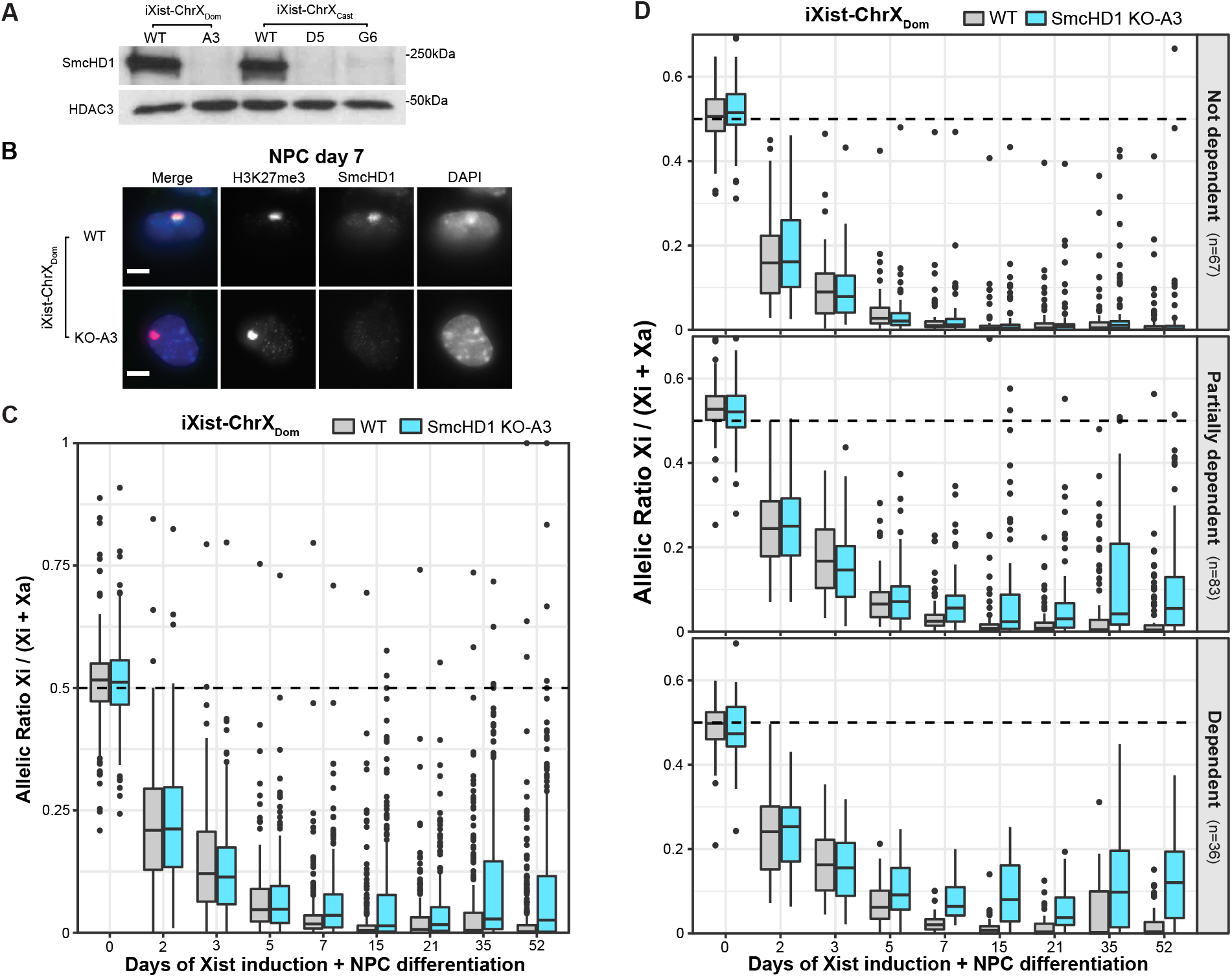
SmcHD1-dependent genes fail to establish complete silencing in SmcHD1 KO A) Western blot demonstrating the absence of SmcHD1 protein in iXist-ChrX KO clones. HDAC3 is included as a loading control. Molecular weight markers are shown on the right. B) Immunofluorescence analysis of SmcHD1 in WT and iXist-ChrX_Dom_ KO clone A3 at NPC differentiation day 7, illustrating the absence of SmcHD1 over the Xi (as marked by an enriched domain of the Polycomb modification H3K27me3). H3K27me3 is shown in magenta, SmcHD1 in cyan, and DNA in blue (DAPI) in the merged image on the left. Scale bar is 5 *µ*m. C) Boxplots summarizing allelic ChrRNA-seq analysis of X-linked gene expression in iXist-ChrX_Dom_ WT and SmcHD1 KO clone A3 (n=246 genes). Boxes of days 0 to 7 of Xist induction are averaged from two replicate timecourse experiments. NPC boxes (days 15 - 52) show individual samples (two collected per NPC timecourse). D) Boxplots summarizing allelic ChrRNA-seq analysis of X-linked gene expression in iXist-ChrX_Dom_ WT and SmcHD1 KO clone A3, with subsets of genes separated by SmcHD1 dependence categories (see Figure S5C). Boxes of days 0 to 7 of Xist induction are averaged from two replicate timecourse experiments. NPC boxes (days 15 - 52) present individual samples, two collected per NPC timecourse.

We went on to analyse Xi gene silencing in SmcHD1 KO mESCs using an extended timecourse of NPC differentiation. Data for a single iXist-ChrX_Dom_- and two iXist-ChrX_Cast_-derived SmcHD1 KO cell lines alongside wild-type controls are presented in Figure 6C and Figure S6D. At early timepoints, up to 5-7 days of differentiation, silencing in SmcHD1 KO cells proceeded with a similar trajectory to that seen in wild-type cells. However, at later timepoints Xi gene silencing plateaued in SmcHD1 KO cells. To investigate this observation further, we analysed silencing for the different SmcHD1 dependency groups as defined above. As shown in Figure 6D, silencing of SmcHD1-‘not dependent’ genes proceeded to completion, whereas both SmcHD1-dependent and partially dependent genes retain some activity on Xi at all stages of NPC differentiation. Similar results were obtained for SmcHD1 KO iXist-ChrX_Cast_ mESCs (Figure S6E). Notably, differences between WT and KO for SmcHD1-dependent and partially dependent genes become apparent at day 5 of NPC differentiation (Figure 6A,B lower panels), which correlates with the time of SmcHD1 recruitment to the inactive X chromosome (Figure 5A,B). Together these results demonstrate that SmcHD1 contributes to the establishment of silencing at a specific subset of genes during Xist-mediated chromosome silencing in a differentiation-dependent manner, and accordingly that completion of the X inactivation process occurs only in cells that have transitioned away from the pluripotent state.

## Discussion

Our findings reinforce the view that Xist-mediated gene silencing is largely attributable to chromatin modification by the SPEN and Polycomb pathways, but also highlight that the two pathways function in parallel. It is important to stress that this is not due to regulation of mutually exclusive gene subsets but rather to a varying contribution to silencing on a gene-by-gene basis. Genes more affected by disruption of the Polycomb system are generally expressed at a lower level in mESCs and are located in chromatin environments with higher initial H3K27me3 levels, indicative of proximity to sites where the Polycomb system is targeted independent of Xist RNA expression. In contrast, genes strongly dependent on the SPEN system tend to be more highly expressed in mESCs (Nesterova et al., 2019), which is in line with recent evidence that SPEN is rapidly recruited to active promoters and enhancers of X-linked genes upon Xist expression (Dossin et al., 2020). That SPEN and Polycomb function in parallel is supported by the observations that SPOC-independent silencing persists in mESCs differentiated into NPCs, and that complete loss of silencing occurs in SPEN^SPOCmut^ mESCs following either depletion of PCGF3/5 or deletion of the Xist B/C-repeat, the Polycomb complex subunits and Xist RNA region required for Xist-mediated Polycomb recruitment respectively. A caveat in reaching these conclusions is that both SPEN^SPOCmut^ and Polycomb mutations have minor effects on Xist RNA behaviour/localization, and that this is increased somewhat in mESCs with both pathways abrogated. However, as noted above, these effects are unlikely to account for the complete loss of silencing that we observe in the latter scenario.

Recent work has shown that the catalytic activity of PRC1 complexes, specifically deposition of H2AK119ub1, is essential to the maintenance of Polycomb target gene repression in mESCs (Blackledge et al., 2020; Tamburri et al., 2020). Previously we reported that the effects of Polycomb on Xist-mediated silencing are largely attributable to PRC1 activity (Nesterova et al., 2019), and it follows that this is likely linked to H2AK119ub1 deposition over Xi. However, exactly how H2AK119ub1 impacts on Xist-mediated silencing remains to be established. Possible mechanisms include direct effects on chromatin structure, for example compaction or transcriptional inhibition, and indirect effects involving reader proteins. In X inactivation, recruitment of the chromosomal protein SmcHD1 provides an example of the latter (Jansz et al., 2018a and see below). However, a contribution of H2AK119ub1 in Xist-mediated silencing is seen prior to SmcHD1 recruitment, demonstrating that other mechanisms, either direct or indirect are also important. Further studies are required to determine if this is linked to direct effects of H2AK119ub1 on chromatin structure/transcription or on other unidentified H2AK119ub1 reader proteins.

Whilst the SPEN and Polycomb pathways are sufficient for initial establishment of Xist-mediated silencing, repression of individual X-linked genes progresses with very different trajectories, ranging from 3-4 hours to several days. In this study we find that for genes that are normally inactivated relatively slowly, cellular differentiation is required to complete the silencing process. Thus, dependence on cellular differentiation for the completion of silencing is apparent for a significant proportion of X-linked genes. Formally this could be attributable to a factor(s) presence of which is limited to differentiated cells, or to different properties of mESCs relative to differentiated derivatives, for example a more rapid cell cycle. Support for the former possibility is exemplified by the chromosomal protein SmcHD1, which is recruited to Xi only in differentiated mESC derivatives. Indeed, our analysis demonstrates a role for SmcHD1 in establishment of silencing, affecting genes with intermediate/slow silencing dynamics. It should be noted that the pathway that elicits Xi SmcHD1 recruitment in differentiated cells is unknown. We speculate that a factor present only in differentiated cells facilitates SmcHD1 recognition of H2AK119ub1. Previous work indicates that this is unlikely to be the loading factor LRIF1, which mediates SmcHD1 recruitment to H3K9me3-modified chromatin via interaction with HP1 proteins, but is dispensable for SmcHD1 localization to Xi in MEFs (Brideau et al., 2015). Similarly, the mechanism of action of SmcHD1 in X inactivation is also poorly understood. Previous work has pointed to functions in chromosome compaction (Nozawa et al., 2013), DNA methylation of CpG islands (Blewitt et al., 2008; Gendrel et al., 2012) and eviction of cohesin/CTCF in the context of formation of Xi specific higher order chromosome structures (Gdula et al., 2019; Jansz et al., 2018b; Wang et al., 2018). Further studies are needed to address this issue, both in terms of SmcHD1 function in establishment and maintenance of Xist-mediated silencing.

In conclusion, our findings provide a comprehensive view of key steps underpinning the establishment of Xist-mediated silencing, as summarized in Figure 7. In a wider context, the role of SmcHD1 in reinforcing gene repression by the Polycomb system specifically in differentiated cells is likely to be relevant at other Polycomb target loci, for example Hox gene clusters, and as such provides an important paradigm for how epigenetic mechanisms contribute to locking gene expression states as cells transition from pluripotency to terminal differentiation.

**Figure 7.**
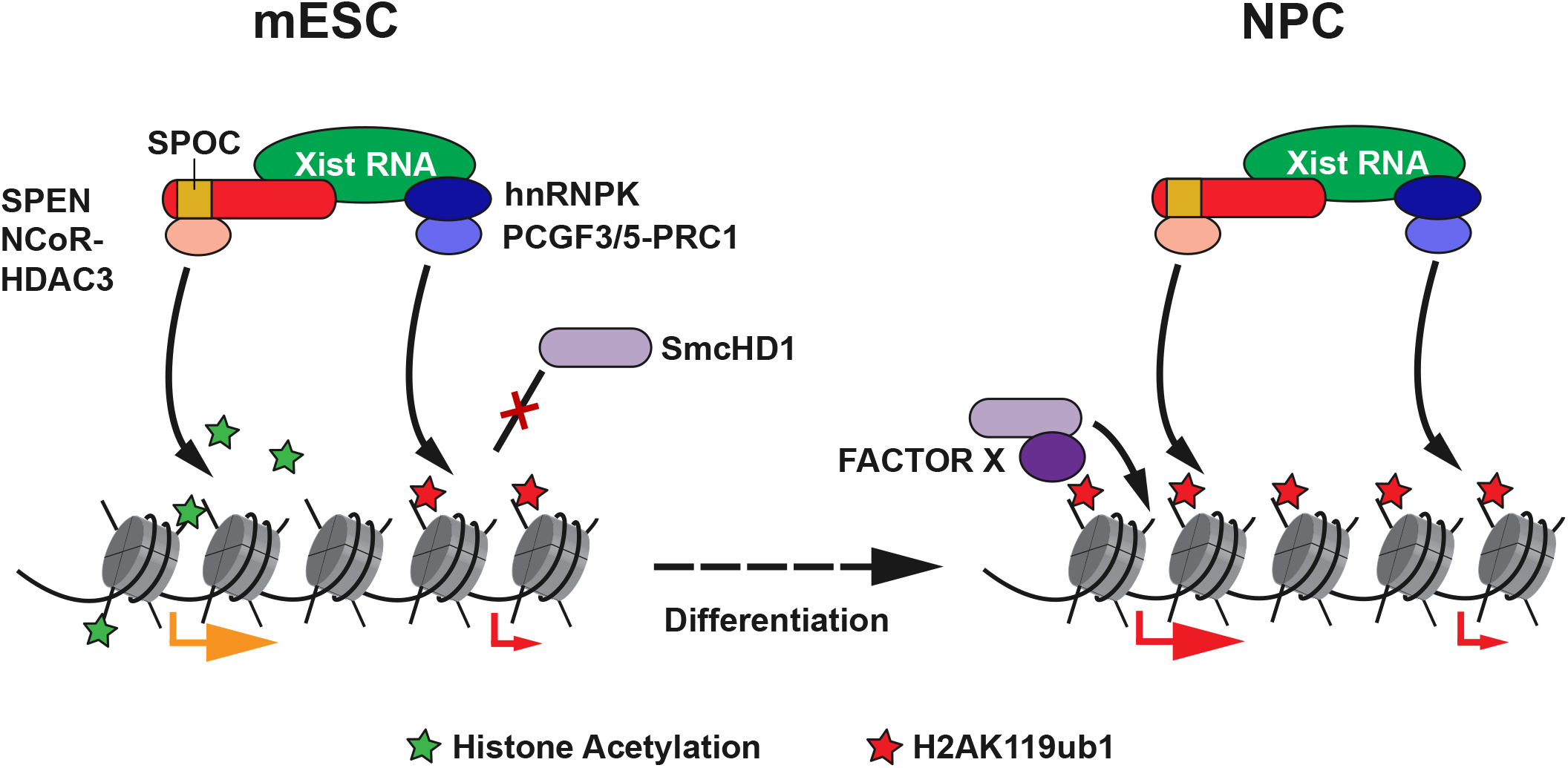
Model illustrating pathways of Xist-mediated gene silencing In undifferentiated mESCs (left), Xist establishes gene silencing principally through pathways downstream of the SPEN SPOC domain, notably NCoR-HDAC3-mediated removal of histone acetylation. The Polycomb system, which is recruited by Xist via hnRNPK-PCGF3/5-PRC1, directs widespread H2AK119ub1 deposition over Xi, and functions in parallel with the SPEN pathway to effect SPOC-independent silencing, predominantly at low-expressed genes (indicated with small red arrow). The combined effect of these two pathways is insufficient to completely silence a subset of genes, including many that are highly expressed prior to X inactivation (indicated with large orange arrow). Upon differentiation (right), SmcHD1 is recruited to Xi, dependent on H2AK119ub1 and an unknown differentiation-linked mechanism (herein ‘Factor X’), and effects completion of the silencing process (large red arrow).

## Acknowledgements

We would like to thank members of the Brockdorff lab for critical discussion and suggestions. We would like to thank Anne-Valerie Gendrel and Emilia Dimitrova for advice with NPC differentiation protocols. We are also grateful to Amanda Williams for loading and maintenance of the Illumina NextSeq machine and Oxford Biochemistry IT support for computing server maintenance. This work was funded by Wellcome Trust grants to N.B. (215513) and J.B. (203817).

## Author contributions

Conceptualization, J.B., T.N., N.B.; Methodology, T.N., J.B., G.W., L.R., M.A., N.B.; Formal Analysis, J.B. and G.W.; Investigation, T.N., J.B., G.W., L.R., H.C., E.C., A.K.; Data Curation, J.B., G.W..; Writing – Original Draft, N.B and J.B.; Writing – Review & Editing, N.B., J.B., T.N., L.R., G.W., M.A.; Visualization, J.B. and T.N., Funding Acquisition, N.B., J.B.; Supervision, N.B.

## METHODS

### RESOURCE AVAILABILITY

#### Lead contact

Further information and requests for resources and reagents should be directed to and will be fulfilled by the lead contact, Neil Brockdorff.

#### Materials availability

All oligos, plasmids, and cell lines generated for this study are available for sharing upon reasonable request to the lead contact.

#### Data and code availability

High throughput sequencing data (ChrRNA-seq and ChIP-seq) generated for this study have been deposited to GEO under accession number XXXXX and are publicly available as of the date of publication. Scripts used for analysis are available at [https://github.com/guifengwei] or [https://github.com/joebowness].

Original unprocessed gel images in this manuscript have been deposited to Mendeley Data and are available following this link XXXXX.

Raw and reconstructed 3D-SIM images have been deposited to the Image Data Resource (https://idr.openmicroscopy.org) under accession XXXX. The programs required to run 3D-SIM image processing and analysis are ImageJ (Fiji distribution with SIMcheck), R, and Octave. Details of makefiles and scripts are provided in our previous study (Rodermund et al., 2021).

Any additional information required to reanalyze the data reported in this paper is available from the lead contact upon request.

### EXPERIMENTAL MODEL AND SUBJECT DETAILS

All female (XX) mouse embryonic stem cell (mESC) lines used for this study were originally derived from the parental F1 2-1 line (129/Sv-CAST/EiJ, a gift from J. Gribnau).

mESCs were routinely maintained in Dulbecco’s Modified Eagle Medium (DMEM; Life Technologies) supplemented with 10% foetal calf serum (FCS; ThermoFisher), 2 mM L-glutamine, 0.1 mM non-essential amino acids, 50 μM β-mercaptoethanol, 100 U/mL penicillin / 100 μg/mL streptomycin (all from Life Technologies) and 1000 U/mL LIF (made in-house). mESCs were grown on gelatin-coated plates under standard culture conditions (37°C, 5% CO_2_, humid) atop a ‘feeder’ layer of MitomycinC-inactivated (Merck Life Science) SNLP mouse fibroblasts and passaged upon ∼80% confluency every 2-3 days using TrypLE Express (ThermoFisher) at room temperature.

*Xist* expression was driven by a TetOn promoter induced by addition of 1 µg/mL of doxycycline (Merck Life Science, D9891). Prior to experiments, cells were pre-plated for 30-40 min on gelatinized dishes to allow feeder cells to preferentially attach, with slower-attaching mESCs then taken from suspension and plated on feederless gelatinized dishes to be harvested for further protocols upon confluency (ie. 2-3 days later).

For calibrated ChIP-seq experiments, *Drosophila* S2 (Sg4) cells were grown adhesively at 25 °C in Schneider’s *Drosophila* Medium (Life Technologies) supplemented with 1 x Pen/Strep and 10% heat-inactivated FCS. Cell counting was performed with a Countess 3 Automated Cell Counter (ThermoFisher).

### METHOD DETAILS

#### Molecular cloning of CRISPR/Cas9 plasmids

Single guide RNAs used for generating CRISPR-Cas9-mediated double strand breaks at target loci were designed using the CRISPOR online tool (Concordet and Haeussler, 2018) and are listed in the Key Resources Table. Sequences in bold are those encoding the sgRNA sequences complementary to target sites in the genome. Cloning into sgRNA plasmid vectors was performed using reverse complement oligos and the single-step digestion-ligation Zhang lab protocol (Broad Institute) into the pX459 background (Ran et al., 2013; Addgene plasmid #62988).

Homology vectors for CRISPR-assisted homologous recombination were cloned by Gibson Assembly using oligonucleotides synthesized from Invitrogen. Briefly, 300-500bp homology fragments were amplified by PCR from iXist-ChrX genomic DNA using FastStart High Fidelity enzyme (Merck Life Science). N-terminal FKBP12^F36V^ fragments were originally amplified originally from pLEX_305-N-dTAG (Addgene #91797) (Nabet et al., 2018). Gibson assembly ligation into a restriction-enzyme digested pCAG backbone plasmid was then performed using Gibson Assembly Master Mix (NEB) according to manufacturer’s guidance.

Products from digestion-ligation or Gibson assembly reactions were XL10-Gold ultracompetent bacteria (Agilent). DNA was isolated from bacterial colonies using the Miniprep kit (Qiagen) and confirmed as containing the desired plasmid via Sanger sequencing.

#### Derivation of cell lines

Generation of the parental iXist-ChrX cell lines is described in our previous study (Nesterova et al., 2019) (iXist-ChrX_Dom_ = clone B2, iXist-ChrX_Cas_ = clone C7). SPEN^SPOCmut^ and FKBP12^F36V^-PCGF3/5 mutant cell lines were generated by CRISPR-assisted homologous recombination by co-transfection of cloned homology and Cas9-sgRNA plasmids at a molar ratio of 6:1 (2.5 μg of homology vector, ∼1 μg of sgRNA vector). SmcHD1 KO cells were transfected with 1 μg of each Cas9-sgRNA plasmids for CRISPR/Cas9-mediated mutagenesis. Transfections were performed as follows: 1-1.5 x 10^6^ mESCs were plated into wells of a 6-well plate ∼24 hours prior to transfection. Pen/strep were removed from the growth media ∼3 hours prior to transfection of plasmid vectors using Lifofectamine2000 (ThermoFisher) according to the manufacturer’s protocol. The following day, each well was split into several 10 mm plates at low density and cells were subjected to puromycin selection (∼3 μg/mL) from 48 to 96 hours post-transfection. Following puromycin wash-out, cells were grown under regular mESCs conditions for a further 8-10 days until clonal colonies could be isolated in 96-well plates and positive clones validated and expanded.

Summary details for all mESC lines used in this study, sgRNA sequences and plasmid vectors can be found in the Key Resources Table. Oligonucleotides used for screening and PCR validation of cell lines are also listed in the Key Resources Table.

#### Neural progenitor cell (NPC) differentiation protocol

mESC to NPC differentiation protocols from the literature (Conti et al., 2005; Splinter et al., 2011) were adapted and optimized for iXist-ChrX lines as follows. mESCs were first extensively separated from feeder cells by pre-plating four times for 35-40 min each. 0.5 x 10^6^ mESCs were then plated to gelatin-coated T25 flasks and grown in N2B27 media (50:50 DMEM/F-12:Neurobasal (Gibco) supplemented with 1 X N2 and 1 X B27 (ThermoFisher), 1 mM L-glutamine, 100 μM β-mercaptoethanol, 50 U/mL penicillin / 50 μg/mL streptomycin (all from Life Technologies) with 1 μg/mL doxycycline for continuous Xist induction. On day 7, cells were detached from the base of the flask with Accutase (Merck Life Sciences), and 3 x 10^6^ cells were plated to grow in suspension in 90 mm bacterial petri dishes containing N2B27 + Dox media supplemented with 10 ng/mL EGF and FGF (Peprotech). On day 10, embryoid-body-like cellular aggregates were collected by mild centrifugation (100 g for 2 min) and plated back onto gelatin-coated 90 mm dishes in N2B27 + Dox + FGF/EGF media. When NPC outgrowths reached ∼80% confluency, cells were detached by Accutase treatment and split at 1:3 – 1:4 ratio on gelatinized 90mm petri dishes. For long term differentiated timepoints cells were maintained in N2B27 + Dox + FGF/EGF media and passaged as above. For dTAG13-treated FKBP12^F36V^-PCGF3/5 and combined FKBP12^F36V^-PCGF3/5 + SPEN^SPOCmut^ lines, 100 μM dTAG-13 was added 12 hours prior to initial pre-plating and maintained in the growth media throughout the protocol.

#### Nuclear extraction

Nuclear extracts were made from cell pellets of confluent 90mm dishes (3 x 10^7^ cells, 1 x packed cell volume, PCV). Briefly, cell pellets were washed with PBS then resuspended in 10 x PCV buffer A (10 mM HEPES-KOH pH7.9, 1.5 mM MgCl_2_, 10 mM KCl, with 0.5 mM DTT, with freshly added 0.5 mM phenylmethylsulfonyl fluoride (PMSF) and complete protease inhibitors (Roche)). After 10 min on ice to allow cell swelling, cells were centrifuged (1500 g for 5 min at 4°C) and resuspended in 3 x PCV buffer A + 0.1% NP40 (Merck Life Science). After 10 further min on ice, nuclei were collected by centrifugation (400 g for 5 min at 4°C) then resuspended in 1 x PCV buffer C (250 mM NaCl, 5 mM HEPES-KOH pH 7.9, 26% glycerol, 1.5 mM MgCl_2_, 0.2 mM EDTA-NaOH pH8, with fresh 0.5 mM DTT and protease inhibitors). NaCl was then added dropwise up to a concentration of 350 mM, and the extract was incubated for 1 hour on ice with occasional agitation. After centrifugation (16000 g for 20 min at 4°C) the supernatant was taken as soluble nuclear extract. This was quantified by Bradford’s assay (Bio-Rad) and stored at −80 °C until use.

#### Western blotting

Nuclear extracts were used for Western blot analysis of all proteins shown in this study. Samples were loaded onto home-made polyacrylamide gels and transferred to PVDF membranes using the Trans-blot Turbo (Bio-Rad) “Mixed Mw” setting. Membranes were then blocked for 1 hour at room temperature in 10mL blocking buffer: 100mM Tris pH 7.5, 0.9% NaCl, 0.1% Tween (TBST) and 5% Marvel milk powder. Blots were incubated overnight at 4°C with primary antibody (see Key Resources Table), washed four times with blocking buffer, then incubated with constant agitation at room temperature for 1 hour in secondary antibody of the relevant species conjugated to horseradish peroxidase. After washing twice more in blocking buffer, once in TBST, and once in PBS (10 min each), membranes were developed and visualized using Clarity Western ECL substrate (Bio-Rad).

#### Immunofluorescence

Immunofluorescence was performed as described in our previous study (Nesterova et al, 2019). Cells were grown on gelatinized slides or 13 mm diameter coverslips (VWR) for at least a day prior to experimental fixation. Slides were washed with PBS and then fixed with 2% formaldehyde for 15 min followed by 5 min of permeabilization in 0.4% Triton X-100. All procedures were performed at room temperature (RT). Cells were briefly washed with PBS before blocking with a 0.2% PBS-based solution of fish gelatin for three washes of 5 min each. Primary antibody dilutions were prepared in fish gelatin solution with 5% normal goat serum and added to cells on slides for 2 hours of incubation in a humid chamber at room temperature. Slides were then washed three times in fish gelatin solution. Secondary antibodies were diluted 1:400 in fish gelatin solution and incubated with cells on slides for 1 hour in a humid chamber. After incubation, slides were washed twice with fish gelatin and one time with PBS before mounting using Vectashield mounting medium with DAPI. Excess mounting medium was removed and the coverslips were sealed to slides using nail varnish. Cells were analysed and scored for the presence of Xi domains on an inverted fluorescence Axio Observer Z.1 microscope using a PlanApo ×63 /1.4 NA oil-immersion objective. Images were acquired using AxioVision software.

#### Xist RNA FISH

Cells for each sample were split to grow on gelatin-coated 22mm coverslips in wells of 6-well plates and fixed at ∼70% confluency after ∼48 hours. Xist expression was induced for 1 day, and in the case of FKBP12^F36V^ lines dTAG-13 was added 12 hours prior to doxycycline addition. At collection, cells on coverslips were washed once with PBS, fixed in the 6-well plate with 3% formaldehyde pH 7 for 10 min, then washed once with PBS, twice with PBST.5 (0.05% Tween20 in PBS), and transferred into a new 6-well dish for permeabilization in 0.2% Triton X-100 in PBS for 10 min at RT. After three further PBST.5 washes, cells on cover slips were subjected to ethanol dehydration by an initial incubation with 70% EtOH (for 30 min at RT), then progressive exchanges to 80%, 90% and finally 100% EtOH. Xist FISH probe was prepared, starting on the previous day, from an 18 kb cloned cDNA (pBS_Xist; Key Resources Table) spanning the whole Xist transcript using a nick translation kit (Abbott Molecular). The FISH hybridisation mix consisted of: 3 μL Texas Red-labelled Xist probe (∼50 ng DNA), 1 μL 10 mg/mL Salmon Sperm DNA, 0.4 μL 3 M NaOAc and 3 volumes of 100% EtOH per sample. This was precipitated by centrifugation (20,000 g for 20 min at 4°C), washed with 70% EtOH, dried by air or speed vacuum, resuspended in 6 μL deionized formamide (Merck Life Science) per hybridisation, then incubated in a shaker (1400 rpm) at 42°C for at least 30 min. 2X hybridisation buffer (4X SSC, 20% dextran sulphate, 2 mg/mL BSA (NEB), 1/10 volume nuclease free water and 1/10 volume vanadyl-ribonucleoside complex (VRC; pre-warmed at 65°C for 5 min before use)) was denatured at 75°C for 5 min, placed back on ice to cool, then mixed with hybridization mix. Each coverslip was hybridized with 30 μL probe/hybridisation mix in a humid box at 37°C overnight. The next day, coverslips were washed 3 times for 5 min at 42°C with pre-warmed 50% formamide/2X saline-sodium citrate buffer (1/10 20X SSC in PBST.5), then subjected to further washes (3 x 2XSSC, 1 x PBST.5, 1 x PBS, each for 5 min using a 42°C hot plate) before being mounted with VECTASHIELD with DAPI (Vector Labs) onto Superfrost Plus microscopy slides (VWR). Slides were dried and sealed using clear nail polish prior to imaging.

5-10 images (20-40 cells per image) were acquired with AxioVision software on an inverted fluorescence Axio Observer Z.1 microscope (Zeiss) using a PlanApo ×63/1.4 NA oil-immersion objective. Images for all lines were scored together, blinded, for the presence or absence of a noticeable Xist RNA domain.

#### Xist RNA FISH for 3D-SIM

Cells were grown on 18×18 mm No. 1.5H precision cover slips (Marienfield) for 3D-SIM microscopy and treated as described above. Coverslips were washed twice with PBS and fixed using 3% formaldehyde (pH 7) in PBS for 10 min at room temperature. A stepwise exchange of PBST.5 was carried out, before cells were permeabilized using 0.2% Triton X-100 PBS for 10 min at room temperature, then blocked for 30 min (2% BSA / 0.5% fish skin gelatine / PBST.5, 2U/µL RNAsin Plus (Promega) at room temperature). Coverslips were washed twice in PBST.5, then once in 2x SSC before an overnight incubation at 37 °C with FISH probe/hybridization buffer (prepared as above) in a humid chamber. The following day coverslips were washed with 2x SSC as detailed above, then stained with 2µg / ml DAPI in PBST.5 for 10 min at room temperature. Coverslips were washed again with PBS, then milliQ water before mounting as above but using Vectashield mounting medium (no DAPI) and imaged within a week using the DeltaVision OMX V3 Blaze system (GE Healthcare).

#### 3D-SIM

##### Acquisition

3D-SIM imaging was performed on a DeltaVision OMX V3 Blaze system (GE Healthcare) equipped with a 60x/1.42 NA Plan Apo oil immersion objective (Olympus), pco.edge 5.5 sCMOS cameras (PCO), and 405, 488, 593 and 640 nm lasers. Image stacks were acquired with a z-distance of 125 nm and with 15 raw images per plane (5 phases, 3 angles). Spherical aberration after reconstruction was reduced by using immersion oil of different refractive indices (RIs) matched to respective optical transfer functions (OTFs). Here, immersion oil with an RI of 1.514 was used for the sample acquisition and matched to OTFs generated using immersion oil of RI 1.512 for the blue, and 1.514 for the red channel. OTFs were acquired using 170 nm diameter blue emitting PS-Speck beads and 100 nm diameter green and red emitting FluoSphere beads (Thermo Fisher Scientific).

##### Reconstruction

The raw data was computationally reconstructed with softWoRx 6.5.2 (GE Healthcare) using channel-specific OTFs and Wiener filter settings of 0.005. A lateral (x-y) resolution of approximately 120 nm and an axial (z) resolution of approximately 320 nm was achieved (Miron et al., 2020). All data underwent assessment via SIMcheck (Ball et al., 2015) to determine image quality via analysis of modulation contrast to noise ratio (MCNR), spherical aberration mismatch, reconstructed Fourier plot and reconstructed intensity histogram values. Reconstructed 32-bit 3D-SIM datasets were thresholded to the stack modal intensity value and converted to 16-bit composite z-stacks to discard negative intensity values using SIMcheck’s “threshold and 16-bit conversion” utility and MCNR maps were generated using the “raw data modulation contrast” tool of SIMcheck. To eliminate false positive signals from reconstructed noise, we applied a modulation contrast filtering using an adapted in-house Fiji script (Rodermund et al., 2021; Schindelin et al., 2012). Here, all pixels in the reconstructed dataset where the corresponding MCNR values in the raw data map fall below an empirically chosen threshold of 4.0 are set to zero intensity. Thereafter, the resulting ‘masked’ reconstructed dataset is blurred with a Gaussian filter with 0.8 pixel radius (xy) to smoothen hard edges.

##### Alignment

Color channels were registered in 3D with the open-source software Chromagnon 0.85 (Matsuda et al., 2018) determining alignment parameter (x,y,z-translation, x,y,z-magnification, and z-rotation) from a 3D-SIM dataset acquired on the date of image acquisition of multicolor-detected 5-ethenyl-2’-deoxyuridine (EdU) pulse replication labelled C127 mouse cells serving as biological 3D alignment calibration sample (Rodermund et al., 2021).

##### Image analysis workflow

Reconstructed 3D-SIM image stacks were pre-processed and subjected to modulation contrast filtering as described above (Rodermund et al., 2021). Thereafter, lateral color channel alignment was performed as described above. The resulting images were used as representative images of whole nuclei. For further analysis however, the DAPI channel was discarded, and Xist territories were cropped manually using Fiji to exclude signal from neighboring cells. The cropped dimensions were later used to define Xist territory volume in all different cell types.

The processed 3D-SIM image files were analyzed using an in-house adapted makefile script for masking of the signal and centroid determination by watershed algorithm (Rodermund et al., 2021). The output data was used to determine the number of Xist foci in all different cell types. Additionally, localization phenotypes observed in the different cell lines and conditions were scored by eye. Xist RNA territories were scored as either “localized”, “slightly dispersed” or “fully dispersed” based on the fully processed 3D-SIM images.

##### Statistical analysis

For the comparison of two independent large datasets, statistical significance was determined by conducting an unpaired two-sample Wilcoxon test using R. This determines whether it is equally likely that a randomly selected value from one dataset will be less than or greater than a randomly chosen value from another population, making it ideal to determine whether two large datasets have the same distribution. Hence, the unpaired two-sample Wilcoxon test was used as a non-parametric alternative to the unpaired t-test to determine statistical significance.

#### Chromatin RNA-seq

Between 5 x 10^6^ (NPC) and 3 x 10^7^ (mESC) cells were collected from confluent 90mm dishes, washed once with PBS, then snap-frozen and stored at −80°C. Chromatin extraction was performed as follows: Cell pellets were lysed on ice for 5 min in RLB (10 mM Tris pH7.5, 10 mM KCl, 1.5 mM MgCl_2_, and 0.1% NP40). Nuclei were then purified by centrifugation through 24% sucrose/RLB (2800 g for 10 min at 4°C), resuspended in NUN1 (20 mM Tris pH7.5, 75 mM NaCl, 0.5 mM EDTA, 50% glycerol, 0.1 mM DTT), and then lysed by gradual addition of an equal volume NUN2 (20 mM HEPES pH 7.9, 300 mM NaCl, 7.5 mM MgCl_2_, 0.2 mM EDTA, 1 M Urea, 0.1 mM DTT). After 15 min incubation on ice with occasional vortexing, the chromatin fraction was isolated as the insoluble pellet after centrifugation (2800 g for 10 min at 4°C). Chromatin pellets were resuspended in 1mL TRIzol (Invitrogen) and fully homogenenized and solubilized by eventually being passed through a 23-gauge needle 10 times. This was followed by isolation of chromatin-associated RNA through TRIzol/chloroform extraction with isopropanol precipitation. Precipitated RNA pellets were washed twice with 70% ethanol. Final ChrRNA samples were then resuspended in H_2_O, treated with TurboDNAse and measured by Nanodrop (both ThermoFisher). 500 ng – 1 µg of RNA was used for library preparation using the Illumina TruSeq stranded total RNA kit (RS-122-2301).

#### Native ChIP-seq

Calibrated native ChIP-seq was performed largely as described in our previous studies (Nesterova et al., 2019; Rodermund et al., 2021) using buffers supplemented with 5 mM of the deubiquitinase inhibitor N-ethylmaleimide (Merck Life Science) for H2AK119ub1 ChIP. 4 x 10^7^ mESCs and 1 x 10^7^ Drosophila Sg4 Cells (20% cellular spike-in) were carefully counted using a Countess 3 Automated Cell Counter (ThermoFisher) and pooled. Cells were then lysed in RSB (10 mM Tris pH8, 10 mM NaCl, 3 mM MgCl_2_, 0.1% NP40) for 5 min on ice with gentle inversion before nuclei collection by centrifugation (1500 g for 5 min at 4°C). Nuclei were resuspended in 1 mL of RSB + 0.25 M sucrose + 3 mM CaCl_2_, treated with 200U of MNase (Fermentas) for 5 min at 37°C, quenched with 4 µl of 1M EDTA, then centrifuged at 2000 g for 5 min. The supernatant was transferred to a fresh tube as fraction S1. The remaining chromatin pellet was incubated for 1 hour in 300 µl of nucleosome release buffer (10 mM Tris pH7.5, 10 mM NaCl, 0.2 mM EDTA), carefully passed five times through a 27G needle, and then centrifuged at 2000 g for 5 min. The supernatant from this S2 fraction was combined with S1 to make the final soluble chromatin extract. For each ChIP reaction, 100 µl of chromatin was diluted in Native ChIP incubation buffer (10 mM Tris pH 7.5, 70 mM NaCl, 2 mM MgCl_2_, 2 mM EDTA, 0.1% Triton) to 1 mL and incubated with Ab (see Key Resources Table) overnight at 4 °C. Samples were incubated for 1 hour with 40 µl protein A agarose beads pre-blocked in Native ChIP incubation buffer with 1 mg/mL BSA and 1 mg/mL yeast tRNA, then washed a total of four times with Native ChIP wash buffer (20 mM Tris pH 7.5, 2 mM EDTA, 125 mM NaCl, 0.1% Triton-X100) and once with TE pH 7.5. All washes were performed at 4°C. The DNA was eluted from beads by resuspension in elution buffer (1% SDS, 100 mM NaHCO_3_) and shaking at 1000rpm for 30 min at 25°C, and was purified using the ChIP DNA Clean and Concentrator kit (Zymo Research). Enrichment of ChIP DNA at predicted sites for each chromatin modification was confirmed by qPCR using primers given in the Key Resources Table and SensiMix SYBR (Bioline, UK). 25 – 100 ng of ChIP DNA was used for library prep using the NEBNext Ultra II DNA Library Prep Kit with NEBNext Single indices (E7645).

#### Fragment size verification, quantification, and sequencing of NGS libraries

Next-generation sequencing DNA libraries were loaded on a Bioanalyzer 2100 (Agilent) with High Sensitivity DNA chips to verify fragment size distribution between 200-800 bp. Sample libraries were quantified using a Qubit fluorometer (Invitrogen) and, optionally, by qPCR with KAPA Library Quantification DNA standards (Roche) and SensiMix SYBR (Bioline) before being pooled together. 2 x 81-cycle paired-end sequencing was performed using an Illumina NextSeq500 (FC-404-2002).

#### ChrRNA-seq data analysis

The standard ChrRNA-seq data mapping pipeline is reported in our previous study (Nesterova et al., 2019). Briefly, raw fastq files of read pairs were first mapped to rRNA by bowtie2 (v2.3.2; Langmead and Salzberg 2012) and rRNA-mapping reads discarded (typically <2%). The remaining unmapped reads were aligned to an N-masked mm10 genome with STAR (v2.4.2a; Dobin et al., 2013) using parameters: “-outFilterMultimapNmax 1 -outFilterMismatchNmax 4 - alignEndsType EndToEnd”. Aligned reads were assigned to separate files for either the *Cast* or *Dom*/129S genomes by SNPsplit (v0.2.0; Krueger and Andrews, 2016) using the “- paired” parameter and a SNPfile containing the 23,005,850 SNPs between *Cast* and *Dom*/129S genomes (UCSC). Read fragments overlapping genes, for both the ‘unsplit’ and ‘allelic’ files of each sample, were counted by the program featureCounts (Liao et al., 2014) using an annotation file of all transcripts and lncRNAs from NCBI RefSeq and the parameters “-t transcript -g gene id -s 2”.

Principle Component Analysis (PCA) of iXist-ChrX samples was performed using the DESeq R package (Anders and Huber, 2010). A variance stabilizing transformation (VST) was applied to a count matrix containing all samples and the top 500 differentially expressed autosomal genes were taken for calculation of principle components.

Further allelic analysis was performed using R and RStudio on allelic count matrix output files from featureCounts. X-linked genes with at least 10 allelically-assigned fragments (i.e. containing reads that overlap SNPs) in >80% of WT samples were retained for gene silencing analysis. Gene silencing was assessed by calculating the allelic ratio of read counts, given by *Xi/(Xi + Xa)*. An additional filter on the allelic ratio in uninduced mESCs (0.1 < allelic ratio < 0.9) was also applied, as strongly monoallelic genes are likely to be technical artifacts of singular mis-annotated SNPs.

Kinetic modelling of gene silencing dynamics was performed using WT iXist-ChrX samples collected in NPC differentiation timecourse experiments. Exponential model curve fitting was performed using the “nlsLM” function from the “minpack.lm” R package (Elzhov et al., 2016) to a model of the form:

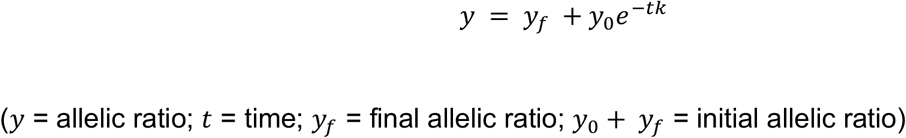

where *y_f_* = 0 is fixed for genes that undergo complete inactivation but is allowed as a parameter for escapees (defined as allelic ratio > 0.1 in mature NPCs). Fitting was done first to the entire dataset in order generate initial parameter estimates. These were then used as inputs for linear regression to fit the model to the silencing trajectory of each gene individually. Model fitting was possible for the vast majority of allelic chrX genes analysed. Silencing halftimes were calculated by the formula:

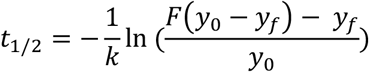

where *k*, and *y_f_* are parameters of the exponential model fit, and *F* = 0.5 (to calculate half of *y*_0_. Halftimes were used to categories X-linked genes into equal classes of fast, intermediate, and slow kinetics of silencing.

For instances where genes are directly categorized or compared by their ‘Initial Expression Level’, this was done using mRNA-seq data from iXist-ChrX_Dom_ mESCs (two replicates averaged together). This data, which contains very few intronic reads, allows for the calculation of a ‘Transcripts per kilobase Million (TPM)’ value for each gene in the count matrix, and hence categorisation of genes into equal groups of low, medium or high expressed. For instances where the relative expression of the same gene (or in the case of Xist, the number of chromatin-associated transcripts) was compared across ChrRNA-seq samples, a simpler RPM (aka CPM; Reads/Counts per Million) transformation of the count matrix was used*.

Summary tables characterizing X-linked genes by all the comparison metrics used in this study are provided in Supplemental File 1.

Plots used to visualize ChrRNA-seq analysis in this study (eg. box, scatter, bar, violin, PCA) were primarily generated using ggplot2 and associated packages in R.

#### Calibrated native ChIP-seq data analysis

For ChIP-seq experiments quantitatively calibrated with *Drosophila* Sg4 cells, raw fastq reads were mapped with bowtie2 (v2.3.2; Langmead and Salzberg, 2012) to the N-masked mm10 genome concatenated with the dm6 *Drosophila* genome. Parameters “–very-sensitive –no-discordant –no-mixed -X 2000” were used and unmapped read pairs were removed. Alignment files were sorted using samtools (Li et al., 2009), PCR duplicates were marked and discarded by the picard-tools “MarkDuplicates” programme, and reads were allelically-assigned using SNPsplit (Krueger and Andrews, 2016). For spike-in calibration, reads mapping to the mm10 and dm6 genomes, in both IP and matched input samples, were counted by samtools. Calibration factors were then calculated according to the derived formula for occupancy ratio (ORi) (Hu et al., 2015) and are provided in Supplemental File 2.

##### Meta-profiles

Meta-profiles from ChIP-seq datasets collected in iXist-ChrX*_Dom_*-derived cell lines were generated from normalized/calibrated bigWig files using the “reference-point” mode of “computeMatrix” in the deeptools suite (Ramírez et al., 2014), followed by the “plotProfile” function. Profiles were centred either on ChrX1 TSSs from gene annotations (UCSC refGene), or published datasets of SUZ12 and RING1B peak locations in mESCs (Fursova et al., 2019).

##### Allelic ChIP-seq analysis

Total and allele-specific alignment (BAM) ChIP and input files were processed into bedGraph format by bedtools “genomeCoverageBed” (Quinlan and Hall, 2010) and calibrated by factors given in Supplemental File 2. The custom Python script *ExtractInfoFrombedGraph_AtBed.py* (https://github.com/guifengwei) was then used to extract values of signal for 250kb windows spanning the 103.5 Mb chrX1 region that can be allelically-analysed. These files were loaded into RStudio for further data processing. Briefly, IP files were first normalized to appropriate input files to calculate enrichment (IP / input) for each window across the chromosome. Line graphs of allelic enrichment were calculated for each sample by subtraction of Xa enrichment from Xi enrichment (Xi – Xa) and are thus ‘internally’ normalized to be more robust to technical variability (eg. in ChIP efficiency) between samples. Data points in boxplots represent allelic enrichment for each window calculated as the ratio of Xi enrichment compared to Xa enrichment (Xi / Xa). ‘Poor mappability’ regions were defined as windows with outlier signal in non-allelic input (+/- 2.5 median absolute deviation). ‘Low allelic’ regions were defined as windows ranking in the lowest 5% of signal in allelic input files. 79 / 414 windows classified as either ‘poor mappability’ or ‘low allelic’ windows are masked by shaded regions in line graphs and excluded from associated boxplots.

**Supplemental File 1** Key information and classification for all X-linked genes amenable to allelic analysis in this study

**Supplemental File 2** Calibration spreadsheet for H2AK119ub1 and H3K27me3 ChIP-seq in FKBP12^F36V^-PCGF3/5

### KEY RESOURCES TABLE

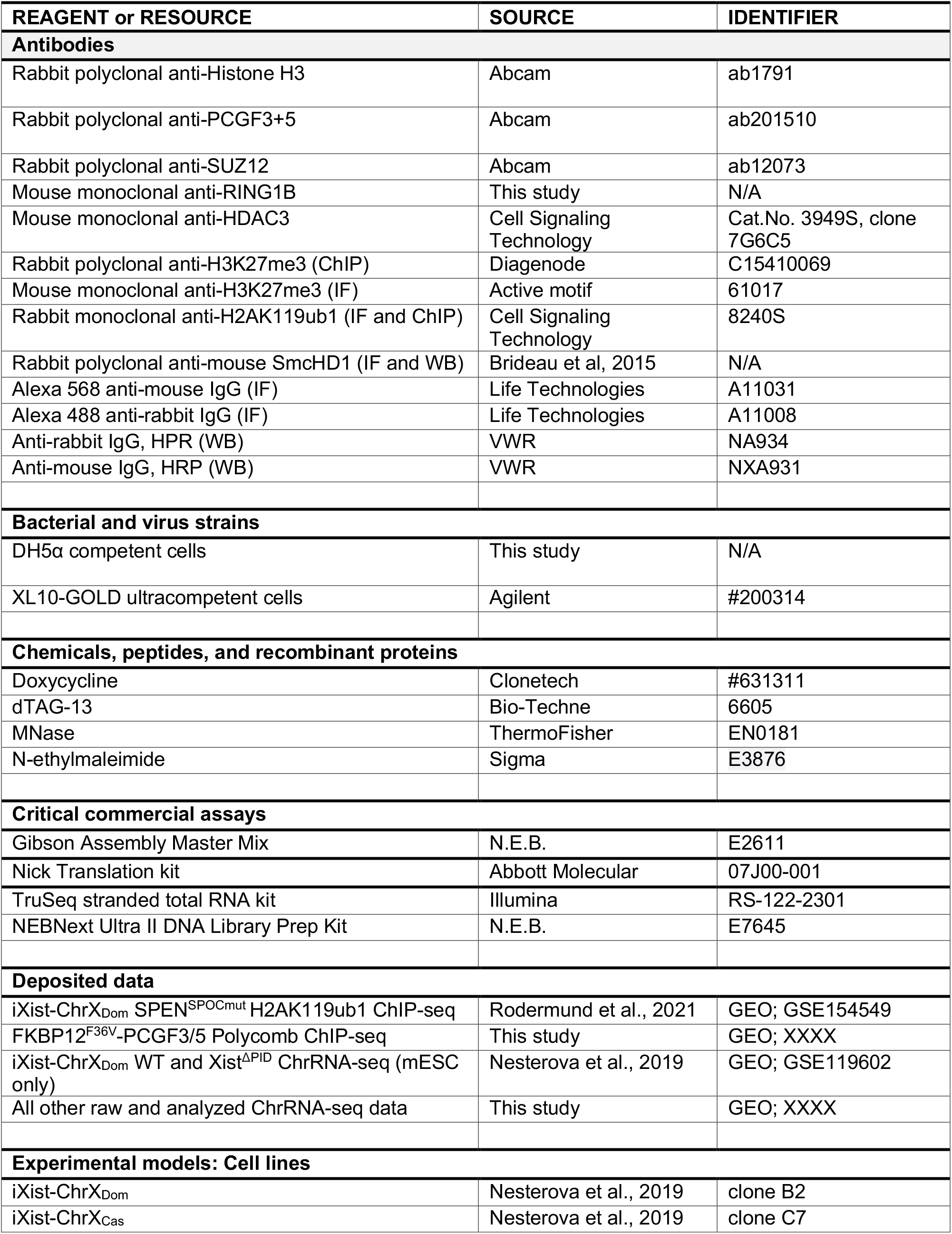
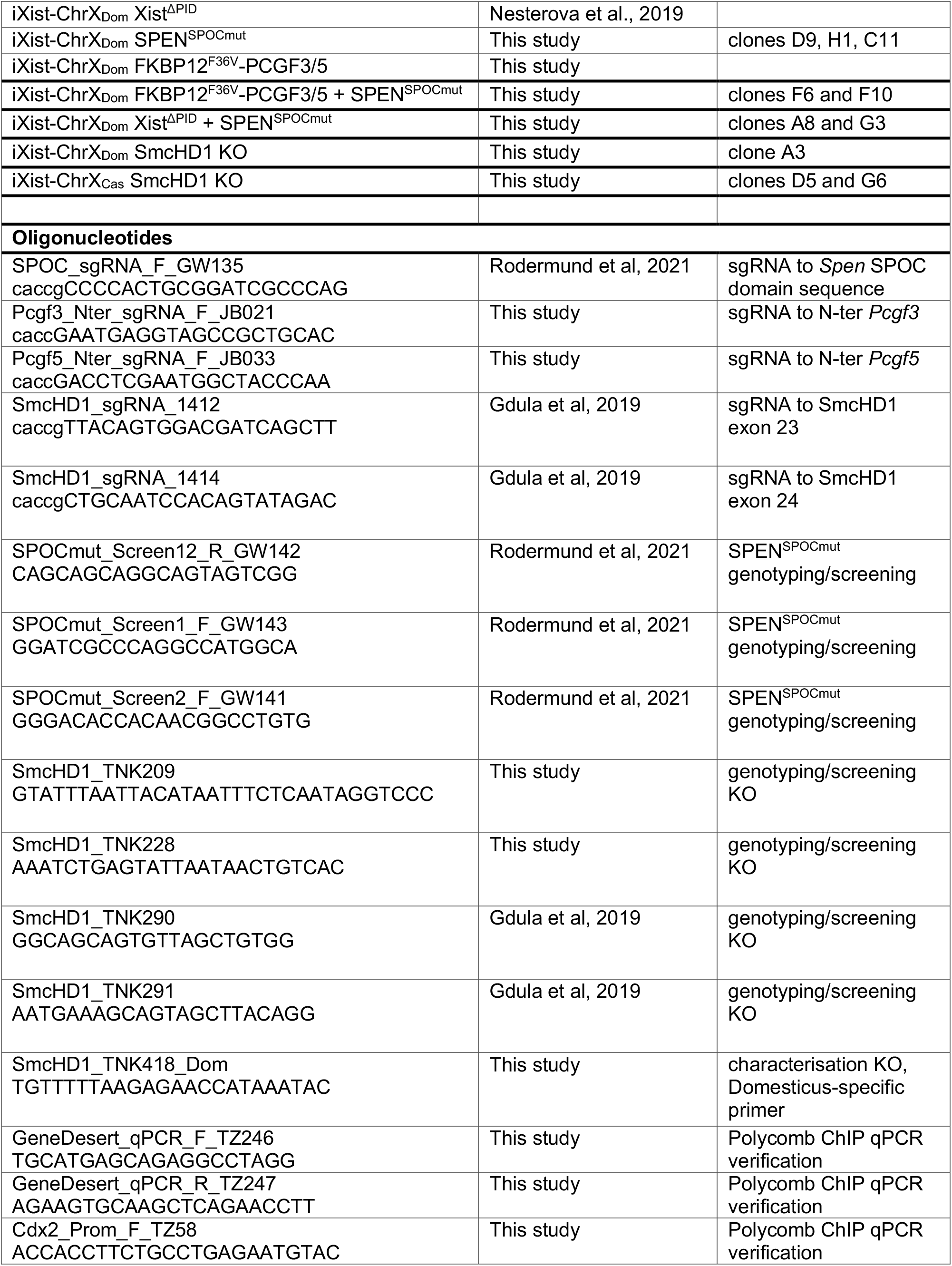
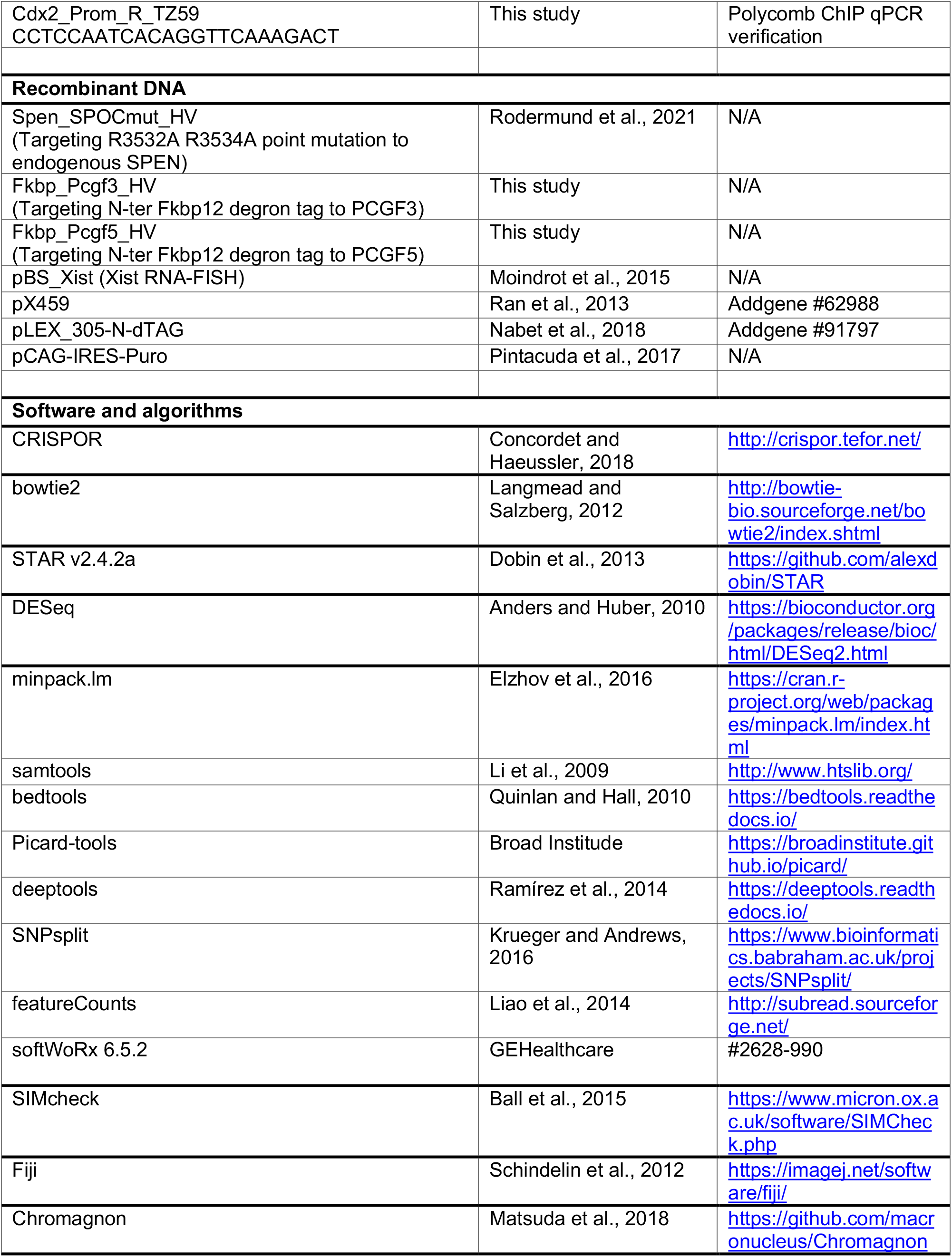

