## Supplemental Figures S1-S6 for "Xist-mediated silencing requires additive functions of SPEN and Polycomb together with differentiation-dependent recruitment of SmcHD1"

This file includes:

**Figures S1-S6**

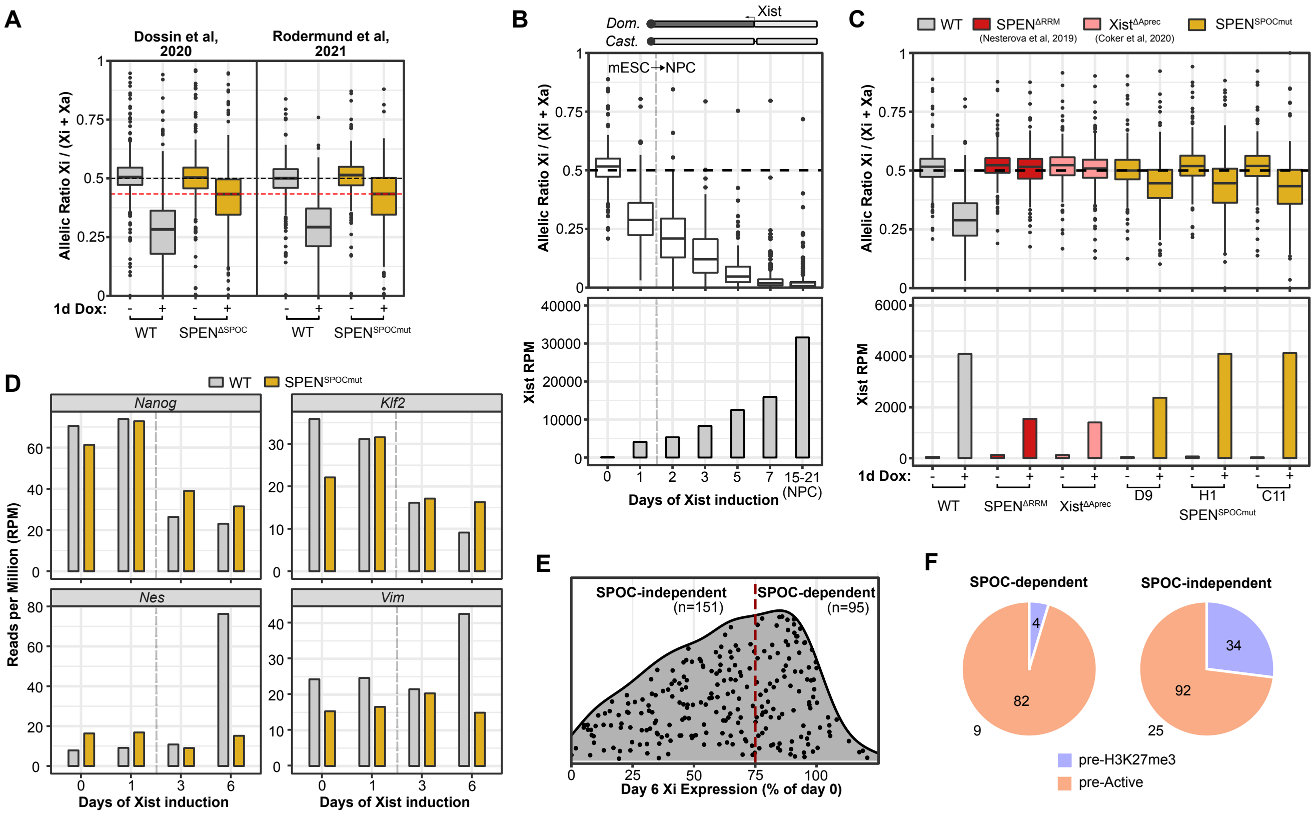


**Figure S1** Characterization of iXist-ChrX$_{Dom}$ SPEN$^{SPOCmut}$. Related to Figure 1.

(A) Boxplots comparing allelic X-linked gene expression between SPEN$^{SPOCmut}$ lines in our previous study (Rodermund et al., 2021) and a larger deletion of the SPEN SPOC domain introduced into a similar model cell line of XX mESCs with inducible Xist (Dossin et al., 2020).

(B) Boxplots summarizing allelic ChrRNA-seq analysis of X-linked gene expression in the iXist-ChrX$_{Dom}$ cell line. Timepoints in mESCs with and without Xist induction (day 1 and day 0 respectively) are shown alongside later timepoints of Xist induction with NPC differentiation. Data from NPC timecourse experiments are averaged (n=2). Relative levels of chromatin-associated Xist RNA for each time point are shown below.

(C) Boxplots summarizing allelic ChrRNA-seq analysis of X-linked gene expression in iXist-ChrX$_{Dom}$ WT, SPEN$^{\Delta RRMmut}$ (Nesterova et al., 2019), Xist$^{\Delta Aprec}$ (Coker et al., 2020) and three independent SPEN$^{SPOCmut}$ clones after 1 day of Xist induction in mESCs. SPEN$^{\Delta RRMmut}$ and Xist$^{\Delta Aprec}$ boxes are averaged from two replicate clones. Relative levels of chromatin-associated Xist RNA for each time point are shown below.

(D) Relative levels of marker gene expression for samples presented in Figure 1A,B.

(E) Methodology used to define a subset of genes that demonstrate greater silencing in SPEN$^{SPOCmut}$ cells. Genes, represented by individual dots, were defined as ‘SPOC-dependent’ if the allelic ratio at day 6 is $>$ 75% of the initial allelic ratio in uninduced day 0 samples.

(F) Pie charts illustrating the proportions of SPOC-independent and SPOC-dependent genes in ‘pre-Active’ and ‘pre-H3K27me3’ gene categories, as defined in (Nesterova et al., 2019). 34 genes were not amenable to ChromHMM analysis in our previous study.

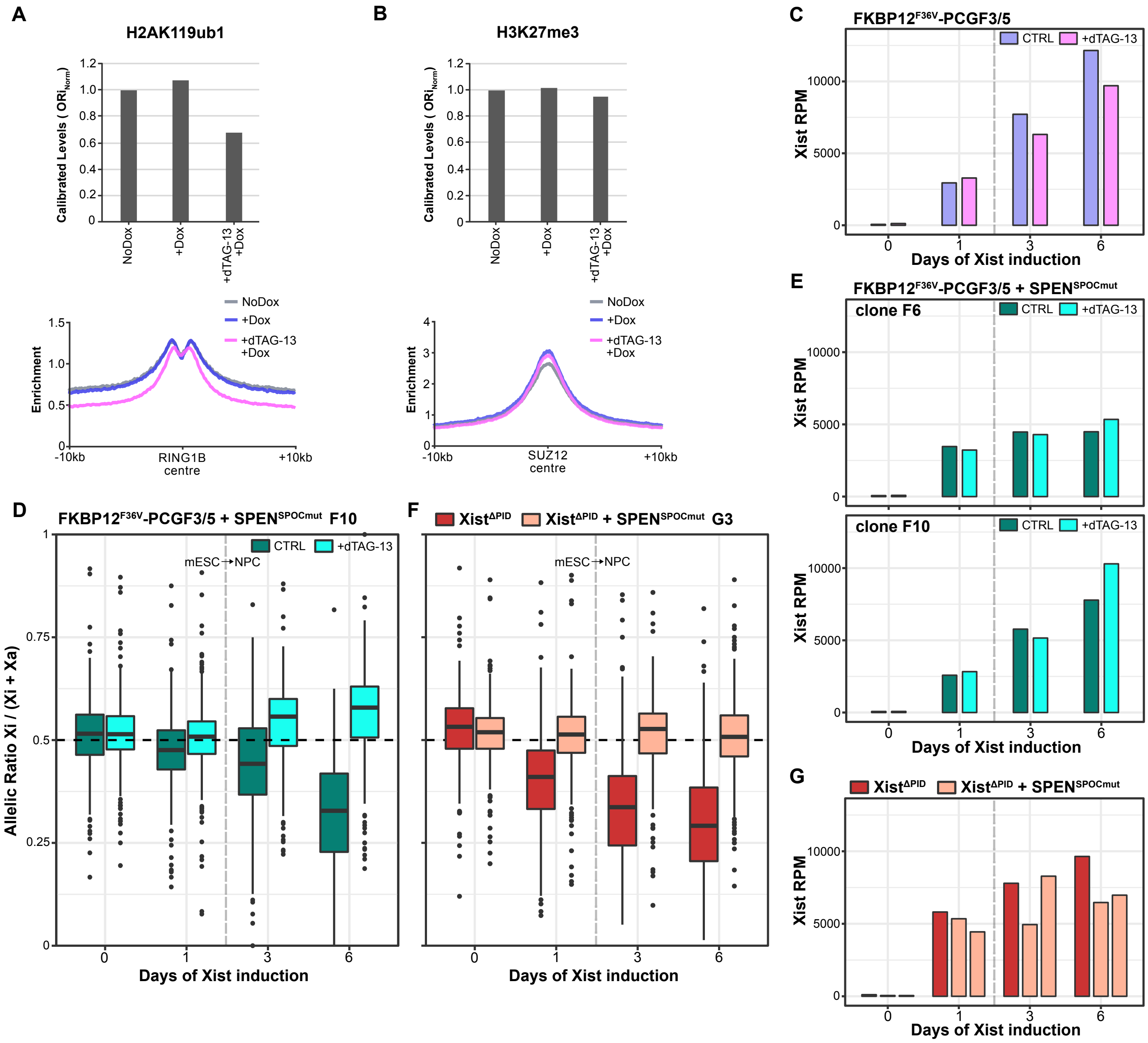


**Figure S2** Characterization of iXist-ChrX$_{Dom}$ FKBP12$^{F36V}$-PCGF3/5 and combined Polycomb + SPEN$^{SPOCmut}$ lines. Related to Figure 2.

(A) (above) Calibrated global levels of H2AK119ub1 from ChIP-seq of FKBP12$^{F36V}$-PCGF3/5 with exogenous spike-in of *Drosophila* cells, averaged from two replicate ChIP-seq experiments (see Methods). (below) Meta-profiles of H2AK119ub1 enrichment centred on PRC1 target regions defined by RING1B ChIP-seq peak centres in mESCs (Fursova et al., 2019).

(B) As (A) for H3K27me3 and PRC2 regions defined by SUZ12 ChIP-seq peak centres in mESCs (Fursova et al., 2019).

(C) Relative levels of chromatin-associated Xist for each sample of the boxplots in Figure 2C.

(D) As Figure 2D for FKBP12$^{F36V}$-PCGF3/5 + SPEN$^{SPOCmut}$ clone G3. The upward skew of allelic ratio for NPC-differentiated samples in this clone is interpreted to be a consequence of selection for cells in the population which have spontaneously eliminated the *Castaneous* X chromosome.

(E) Relative levels of chromatin-associated Xist for each FKBP12$^{F36V}$-PCGF3/5 + SPEN$^{SPOCmut}$

clone, comparing untreated cells and cells treated for 12 hours with dTAG-13 prior to Xist induction cultured in parallel.

(F) As Figure 2E for Xist$^{\Delta PID}$ + SPEN$^{SPOCmut}$ clone G, which is tetraploid and contains two copies of each ChrX allele (see Figure S3D).

(G) Relative levels of chromatin-associated Xist in ChrRNA-seq data sets from Xist$^{\Delta PID}$ and Xist$^{\Delta PID}$ + SPEN$^{SPOCmut}$ clones presented in Figure 2E and Figure S2F.

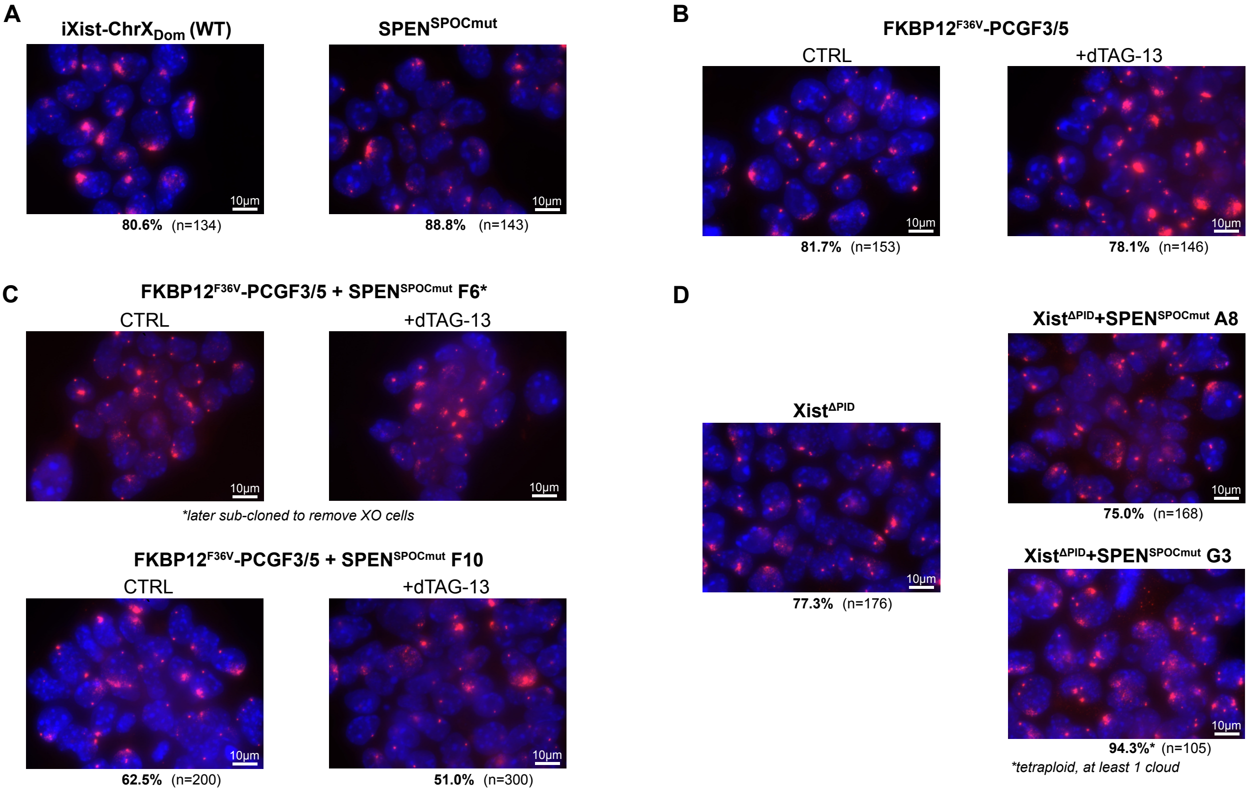


**Figure S3** Efficient Xist induction and cloud formation in SPEN$^{SPOCmut}$ and Polycomb mutant lines. Related to Figure 3.

Xist RNA-FISH images collected from mESCs after 1 day of Xist induction for mutant cell lines presented in Figures 1 and 2 of this study. For the FKBP12$^{F36V}$-PCGF3/5 line and clonal derivatives, dTAG-13 treatment was applied 12 hours prior to Xist induction. The percentage of cells containing Xist clouds is indicated below each representative image. RNA-FISH characterization of Xist induction in FKBP12$^{F36V}$-PCGF3/5 + SPEN$^{SPOCmut}$ clone F6 (C) revealed a sub-population of XO cells with only one Xist/Tsix cloud/foci. This clone was therefore sub-cloned prior to the ChrRNA-seq experiment presented in Figure 2D.

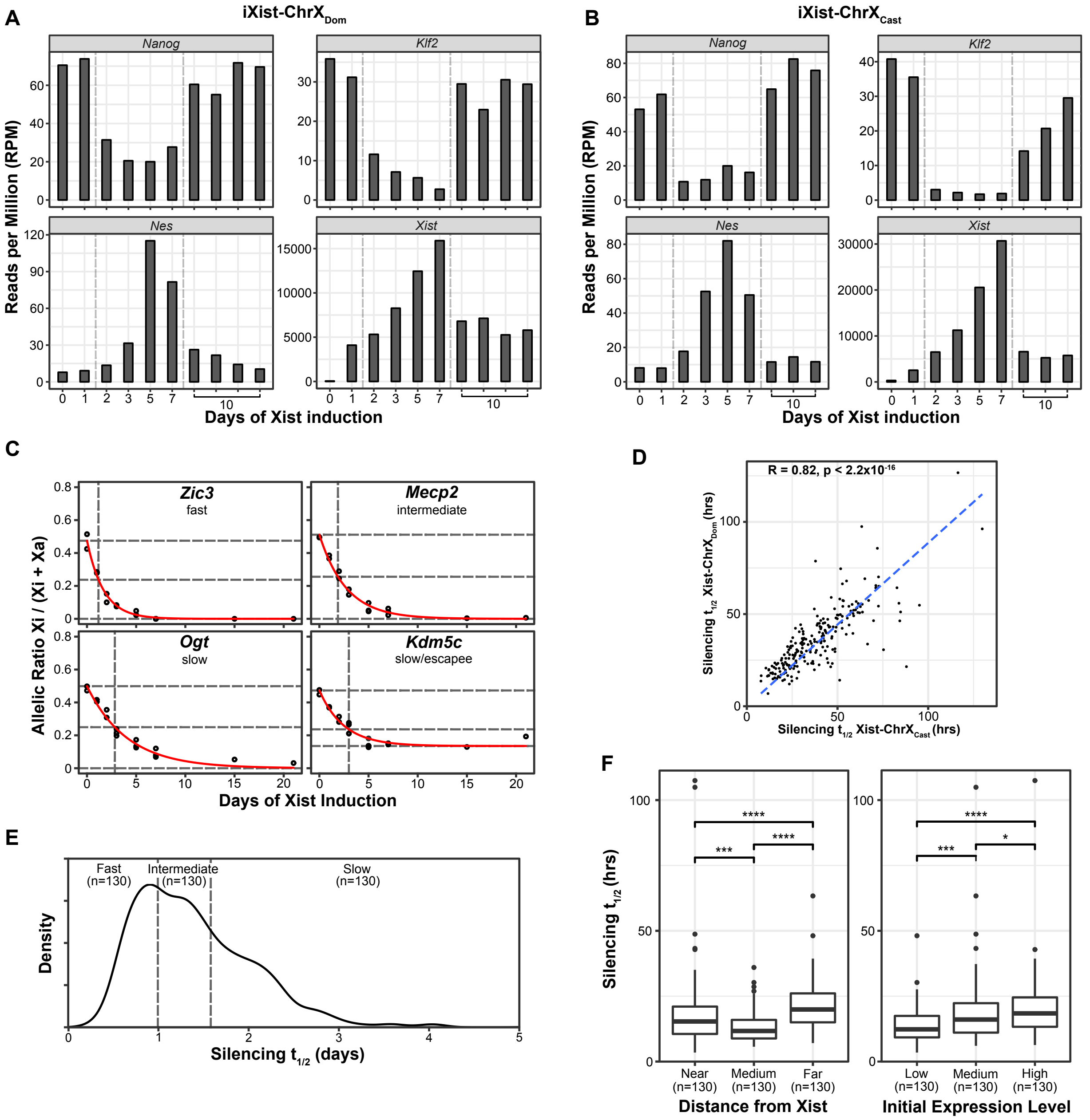

**Figure S4** Modelling of silencing kinetics over a timecourse of Xist induction with NPC differentiation. Related to Figure 4.

(A,B) Relative levels of selected marker gene expression and chromatin-associated Xist for the (A) iXist-ChrX$_{Dom}$ or (B) iXist-ChrX$_{Cast}$ NPC timecourse and undifferentiated day 10 Dox-induced mESC samples shown in with Figure 4A,B.

(C) Silencing trajectories for individual example genes in the iXist-ChrX$_{Cast}$ cell line. *Zic3*, *Mecp2*, *Ogt* are fast, intermediate, and slow silencing respectively. *Kdm5c* is an escapee but is also classified as slow silencing by its initial silencing kinetics. Horizontal lines represent parameters of $y_0$ (initial allelic ratio), 1/2 $y_0$, and $y_f$ (final allelic ratio) respectively, and the vertical line is placed at the calculated $t_{1/2}$ (halftime) for each gene.

(D) Scatter plot comparing calculated silencing halftimes for each gene amenable to allelic analysis of silencing kinetics in both iXist-ChrX$_{Dom}$ and iXist-ChrX$_{Cast}$ lines (n=235). Halftimes are strongly correlated with a Spearman's rank correlation coefficient of R = 0.82.

(E) Density plot of gene silencing halftimes in iXist-ChrX$_{Cast}$, allowing for classification of genes into equal-sized groups of fast, intermediate, and slow silencing genes.

(F) Boxplots comparing silencing halftimes between subsets of genes based on distance from the *Xist* locus (left) or initial expression level in mESCs (right) (see Methods). Significance of individual comparisons is determined by Welch's unequal variances T-test. *, **, ***, **** indicate p below 0.05, 0.01, 0.001 and 0.0001 respectively.

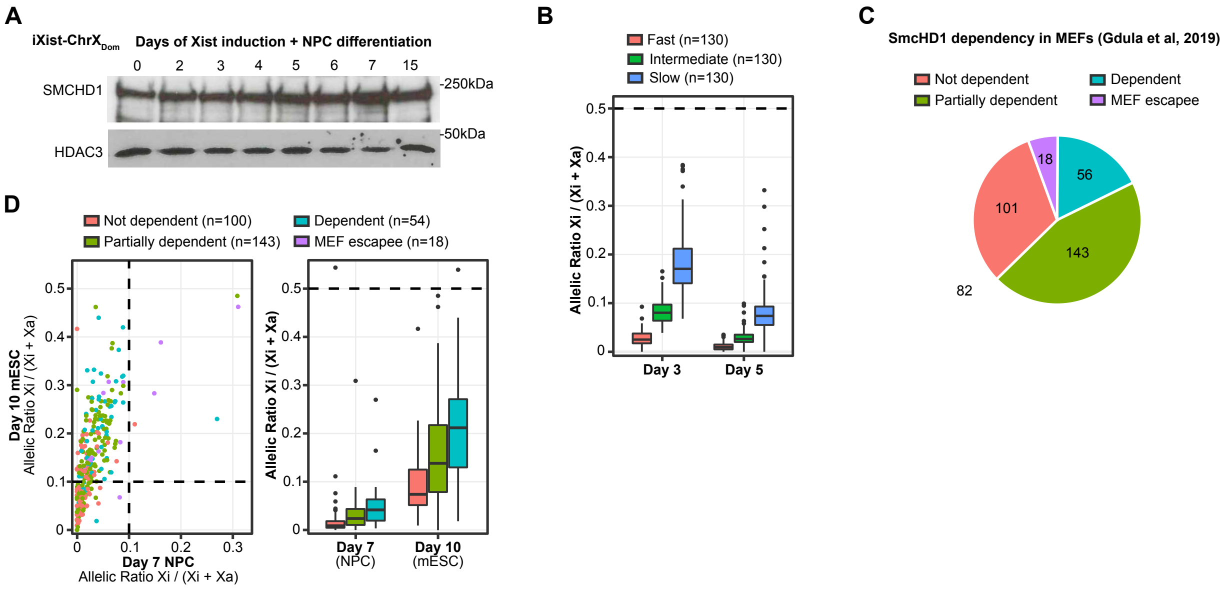


**Figure S5** Slow genes are incompletely silenced at the time window when SmcHD1 is recruited to Xi. Related to Figure 5.

A) Western blot analysis of SmcHD1 protein levels in iXist-ChrX$_{Dom}$ cells during the differentiation timecourse. Note that the amount of SmcHD1 protein present in mESCs (day 0) is comparable to the amount of SmcHD1 detected during differentiation. HDAC3 is included as a loading control. Molecular weight markers are shown on the right.

B) Boxplots summarizing allelic ChrRNA-seq data from iXist-ChrX$_{Cast}$ NPC day 3 and day 5. Boxes compare kinetic classes of genes to demonstrate incomplete silencing of intermediate and slow genes during the time window of SmcHD1 recruitment to Xi.

C) Pie chart depicting the number of genes in each class of SmcHD1 dependency, as defined from ChrRNA-seq analysis of XX mouse embryo fibroblast (MEF) lines derived from SmcHD1 null embryos. 82 genes amenable to analysis in the iXist-ChrX$_{Cast}$ cellular model were not previously categorized due to insufficient expression in MEFs.

D) Scatter plot comparing individual gene allelic ratios between iXist-ChrX$_{Cast}$ undifferentiated mESC day 10 and NPC day 7. Genes are coloured according to SmcHD1 dependency categories. Dashed lines indicate trace allelic ratios of 0.1. Adjacent boxplots compare iXist-ChrX$_{Cast}$ undifferentiated mESC day 10 and NPC day 7 by SmcHD1 dependency categories. MEF escapees are excluded due to insufficient number of genes.

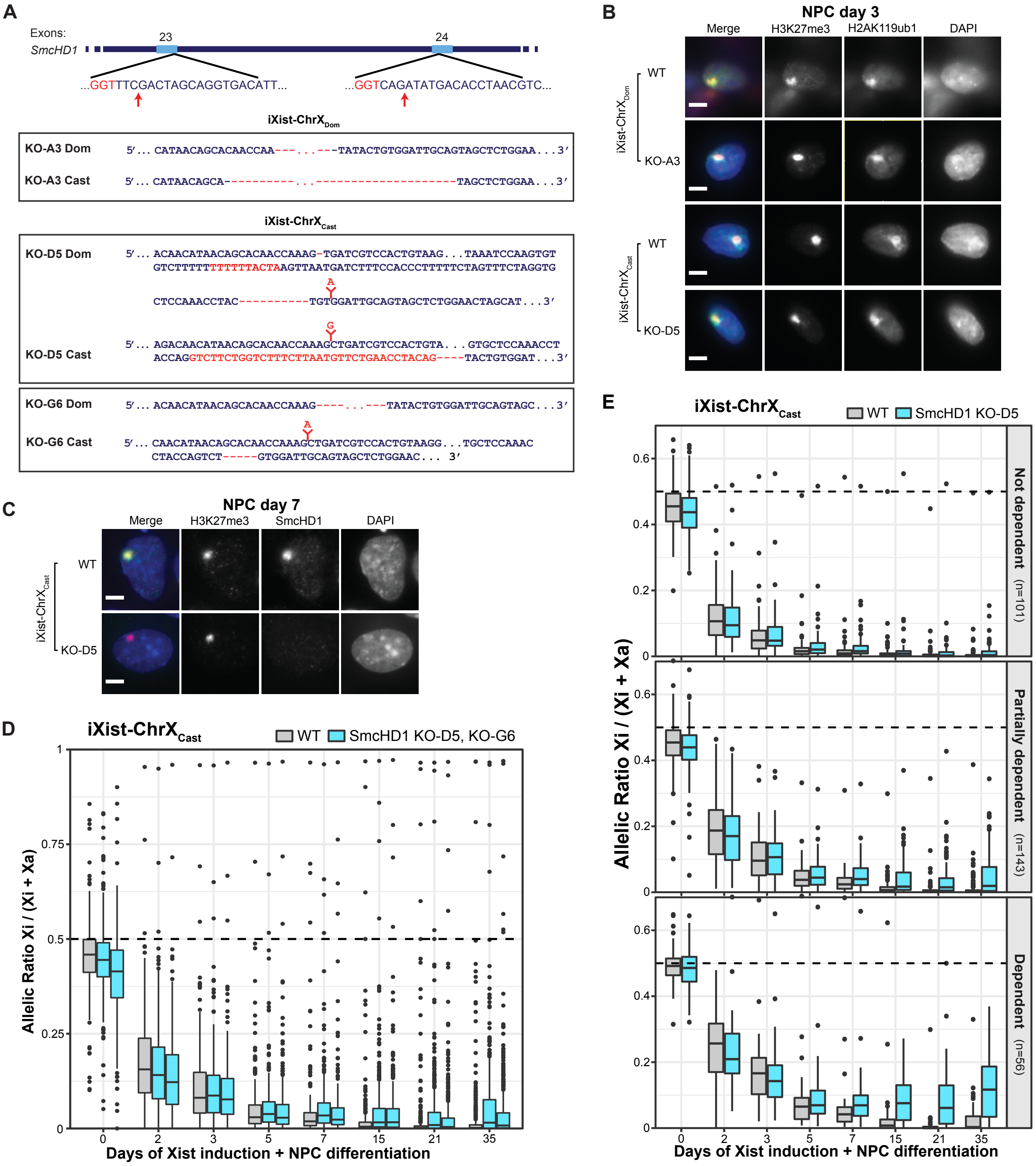


**Figure S6** Characterization of SmcHD1 KO cell lines. Related to Figure 6.

A) Strategy for CRISPR-Cas9-mediated mutagenesis of the *SmcHD1* gene. The positions of guide RNAs relative to *SmcHD1* exons are shown, with PAM sequences (red font) and predicted cleavage sites (red arrows) indicated. Two sgRNAs for exons 23 and 24 were used to generate mutant clones in iXist-ChrX$_{Dom}$ and iXist-ChrX$_{Cast}$. Mutated sequences of *Domesticus* (Dom) and *Castaneous* (Cast) alleles are shown for each KO line. Two independent KO lines were created and analysed for iXist-ChrX$_{Cast}$. Deletions are shown by red dashes and insertions/mutations are shown in red font.

B) Immunofluorescence analysis of Polycomb modifications H3K27me3 and H2AK119ub1 in WT and SmcHD1 KO clones in NPC day 3, confirming Xi domain formation in KO cells. Scale bar is $5\mu$m.

C) Immunofluorescence analysis of SmcHD1 in WT and iXist-ChrX$_{Cast}$ KO clone D5 at NPC differentiation day 7, demonstrating the absence of SmcHD1 over the Xi (as marked by an enriched domain of the Polycomb modification H3K27me3). H3K27me3 is shown in magenta, SmcHD1 in cyan, and DNA in blue (DAPI) in the merged image on the left. Scale bar is $5\mu$m.

D) Boxplots summarizing allelic ChrRNA-seq analysis of X-linked gene expression in iXist-ChrX$_{Cast}$ WT and two independent SmcHD1 KO clones (D5 and G6). Boxes of days 3, 5 and 7 of Xist induction are averaged from three replicate timecourse experiments, day 2 from duplicates, and days 0 and 15 - 35 of SmcHD1 KO from individual samples.

E) Boxplots summarizing allelic ChrRNA-seq analysis of X-linked gene expression in iXist-ChrX$_{Cast}$ WT and SmcHD1 KO (clone D5), with subsets of genes separated by SmcHD1 dependence categories. Boxes of days 3, 5 and 7 of Xist induction are averaged from three replicate timecourse experiments, day 2 from duplicates, and days 0 and 15 - 35 of SmcHD1 KO from individual samples.
